# Admixture has obscured signals of historical hard sweeps in humans

**DOI:** 10.1101/2020.04.01.021006

**Authors:** Yassine Souilmi, Raymond Tobler, Angad Johar, Matthew Williams, Shane T. Grey, Joshua Schmidt, João C. Teixeira, Adam Rohrlach, Jonathan Tuke, Olivia Johnson, Graham Gower, Chris Turney, Murray Cox, Alan Cooper, Christian D. Huber

**Affiliations:** Australian Centre for Ancient DNA, The University of Adelaide, Adelaide, SA 5005, Australia; Transplantation Immunology Group, Immunology Division, Garvan Institute of Medical Research, NSW 2010, Australia; St Vincent’s Clinical School, Faculty of Medicine, UNSW, Darlinghurst, New South Wales, Australia; ARC Centre of Excellence for Mathematical and Statistical Frontiers, The University of Adelaide, Adelaide, South Australia 5005, Australia; Department of Archaeogenetics, Max Planck Institute for the Science of Human History, Jena, Germany; School of Mathematical Sciences, The University of Adelaide, Adelaide, South Australia 5005, Australia; Chronos, University of New South Wales, Sydney, NSW 2052, Australia; Carbon-Cycle Facility and Earth and Sustainability Science Research Centre (ESSRC), University of New South Wales, Sydney, NSW 2052, Australia; Statistics and Bioinformatics Group, School of Fundamental Sciences, Massey University, Palmerston North 4410, New Zealand; South Australian Museum, Adelaide, SA 5005, Australia; BlueSky Genetics, PO 287, Ashton, SA 5137, Australia; Department of Biology, Penn State University, University Park, PA 16802, USA; Melbourne Integrative Genomics, School of BioSciences and School of Mathematics and Statistics, The University of Melbourne, Parkville, VIC 3010, Australia

## Abstract

The role of natural selection in shaping biological diversity is an area of intense interest in modern biology. To date, studies of positive selection have primarily relied upon genomic datasets from contemporary populations, which are susceptible to confounding factors associated with complex and often unknown aspects of population history. In particular, admixture between diverged populations can distort or hide prior selection events in modern genomes, though this process is not explicitly accounted for in most selection studies despite its apparent ubiquity in humans and other species. Through analyses of ancient and modern human genomes, we show that previously reported Holocene-era admixture has masked more than 50 historic hard sweeps in modern European genomes. Our results imply that this canonical mode of selection has likely been underappreciated in the evolutionary history of humans and suggests that our current understanding of the tempo and mode of selection in natural populations may be quite inaccurate.

## Introduction

The rapidly growing availability of genomic datasets provides a powerful resource to address a fundamental question in evolutionary biology, namely the role of natural selection in shaping biological diversity^1^. The proliferation of genomic datasets has been accompanied by parallel developments in statistical methods for uncovering genetic signals of positive selection; however, while the power and precision of these statistical methods has continued to improve, they can be confounded by complex aspects of population history that are not modelled or remain unknown^2^. In particular, previous genomic studies of positive selection typically do not account for past phases of inter-population mixing (i.e. admixture), which can alter genomic signatures of positive selection and either mask these signals or lead to erroneous inferences about the underlying modes of selection^3, 4^.

In the case of humans, a consistent finding has been the apparent paucity of classical ‘hard sweep’ signals in modern genomic datasets (i.e. where a novel beneficial mutation increases to 100% frequency in a population). This has resulted in suggestions that humans adapted to novel environmental and sociocultural pressures through alternate modes of selection – e.g. ‘soft sweeps’ (i.e. where the beneficial allele sits on multiple genetic backgrounds)^5, 6^ or polygenic selection (i.e. subtle frequency shifts across numerous loci with small fitness effects)^7, 8^ – though the evidence for these processes has also been challenged^4, 9–11^. The apparent ubiquity of admixture in human population history^12–14^ suggests instead that the absence of hard sweep signals in modern human genomes may be a consequence of the masking effects of historical admixture events. Moreover, growing evidence suggests that admixture events pervade the history of most natural populations, which may have led to a potentially biased view of the historical role of selection in nature^4, 15^.

### Using ancient human genomes to uncover historical hard sweeps

The recent emergence of population-scale ancient human genomic datasets from Eurasia provides a novel means to search for any hard sweeps that had occurred in the ancestors of modern Europeans, by applying selection scans to genomic datasets that predate known admixture events. Modern Europeans are largely composed of three distinct ancestries – i.e. Western Hunter-Gatherers (WHG), Anatolian Early Farmers (Anatolian EF), and Steppe Pastoralists (see Text S1) – as a result of extensive admixture between these ancestral populations and their descendants that occurred from the Early Holocene to Bronze Age (i.e. ∼12-5 ka;^16^). Accordingly, we reprocessed 1,162 ancient western Eurasian genomes (mostly high-density SNP scans; see Methods) through a uniform bioinformatic pipeline – thereby mitigating potential processing artefacts – and grouped these samples into 18 distinct ancient populations that existed before and after documented Holocene admixture events based on their archaeological and genetic relationships (Figs. 1, S1, S2; Table S1). To ascertain the impact of Holocene admixture on modern European populations, we also analyzed three modern European populations from the 1000 genomes project (1kGP^17^; i.e. Utah residents with Northern and Western European ancestry (CEU); Finnish in Finland (FIN), Toscani in Italy (TSI)), and also included two other non-European 1kGP populations for further comparison (i.e. one East Asian: Han Chinese in Beijing (CHB) and one African: Yoruba in Ibadan (YRI)).

**Fig. 1.**
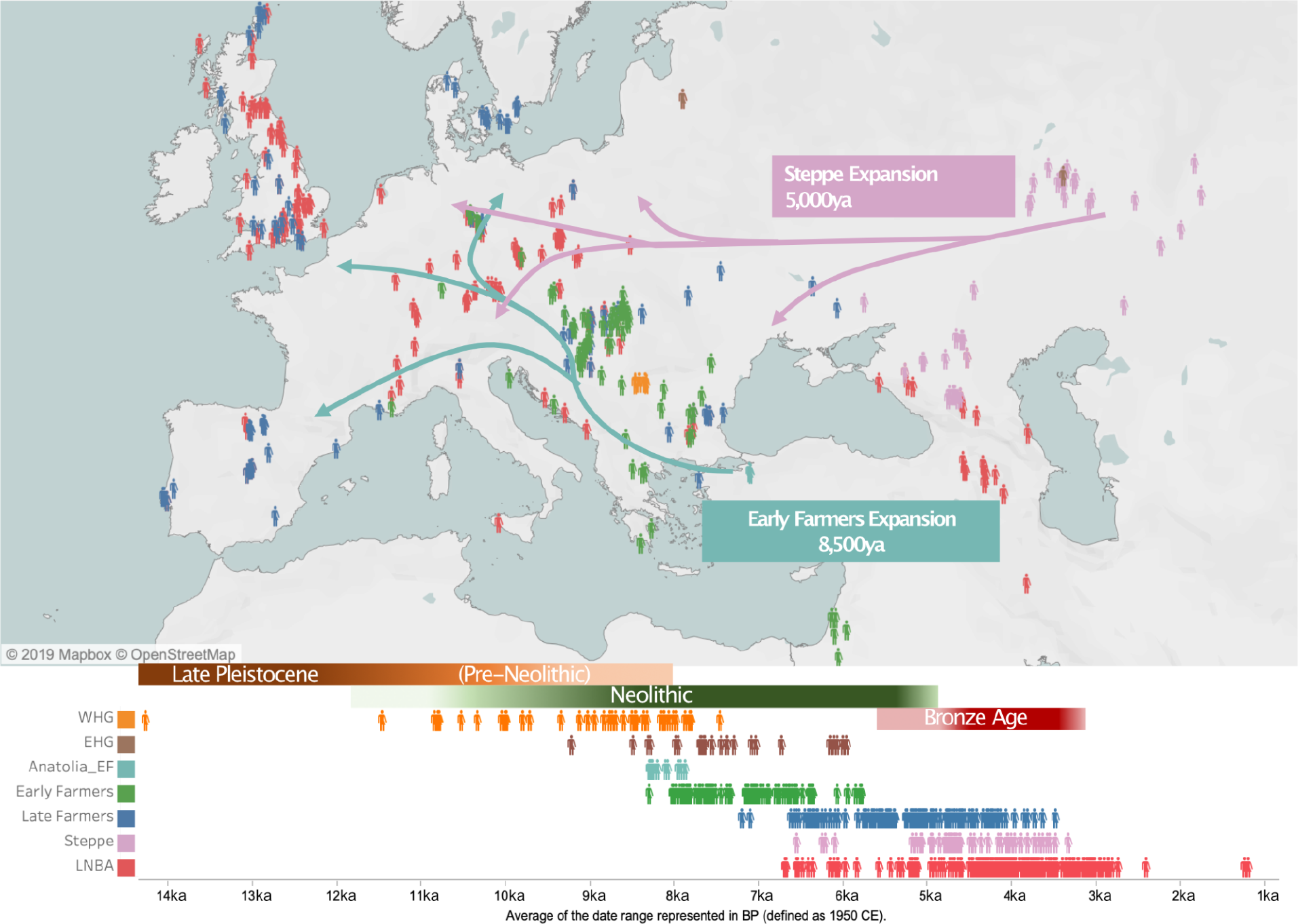
Geographic and temporal distribution of 1,162 ancient Eurasian samples used in this study. Each human symbol represents a sample and the colors indicate different populations classified into broad groupings according to archaeological records of material culture and lifestyle (colors indicated at the bottom right-hand side; see Text S1). The green lines depict the generalized migration route of Anatolian farmers (Anatolia_EF) into Europe ∼8.5ka, where they mixed with Western Hunter-Gatherers (WHG; EHG refers to the contemporaneous Eastern Hunter-Gatherers) to create the Early European Farmers (EF). Similarly, the pink arrows represent the generalized movement of Steppe pastoralists (Steppe; samples east of the Ural Mountains not shown) which resulted in admixture with Late European Farmers (LF) ∼5ka, giving rise to Late Neolithic and Bronze Age (LNBA) societies.

We used SweepFinder2 (SF2)^18^ to scan the genomes of each ancient and modern population for regions exhibiting distorted allele frequency patterns characteristic of a fixed hard sweep (i.e. an excess of low- and high-frequency alleles^19^; also see Text S2). By contrasting each window against site frequency spectrum (SFS) patterns from the overall genomic background, SF2 tests explicitly for the signature of hard selective sweeps while controlling for demographic history and population structure, which often cause false positives in such tests^2, 18^. A set of candidate sweep regions was determined by first assigning SF2 scores to a set of ∼19,000 annotated human genes, and subsequently aggregating neighboring outlier genes into single sweep regions (Fig. S3; see Methods). Importantly, testing our sweep detection pipeline on simulated genomic datasets modelled upon Eurasian demographic history confirmed that it is robust to the impacts of substantial demographic bottlenecks – including strong bottlenecks associated with the founding Eurasian and subsequent WHG populations (successive 12-fold and 6-fold population size reductions, respectively; see Fig. 2A) – and also to missing data, ascertainment bias, and alignment error (Fig. 2B and Figs. S4-S7, SI Methods).

**Fig. 2.**
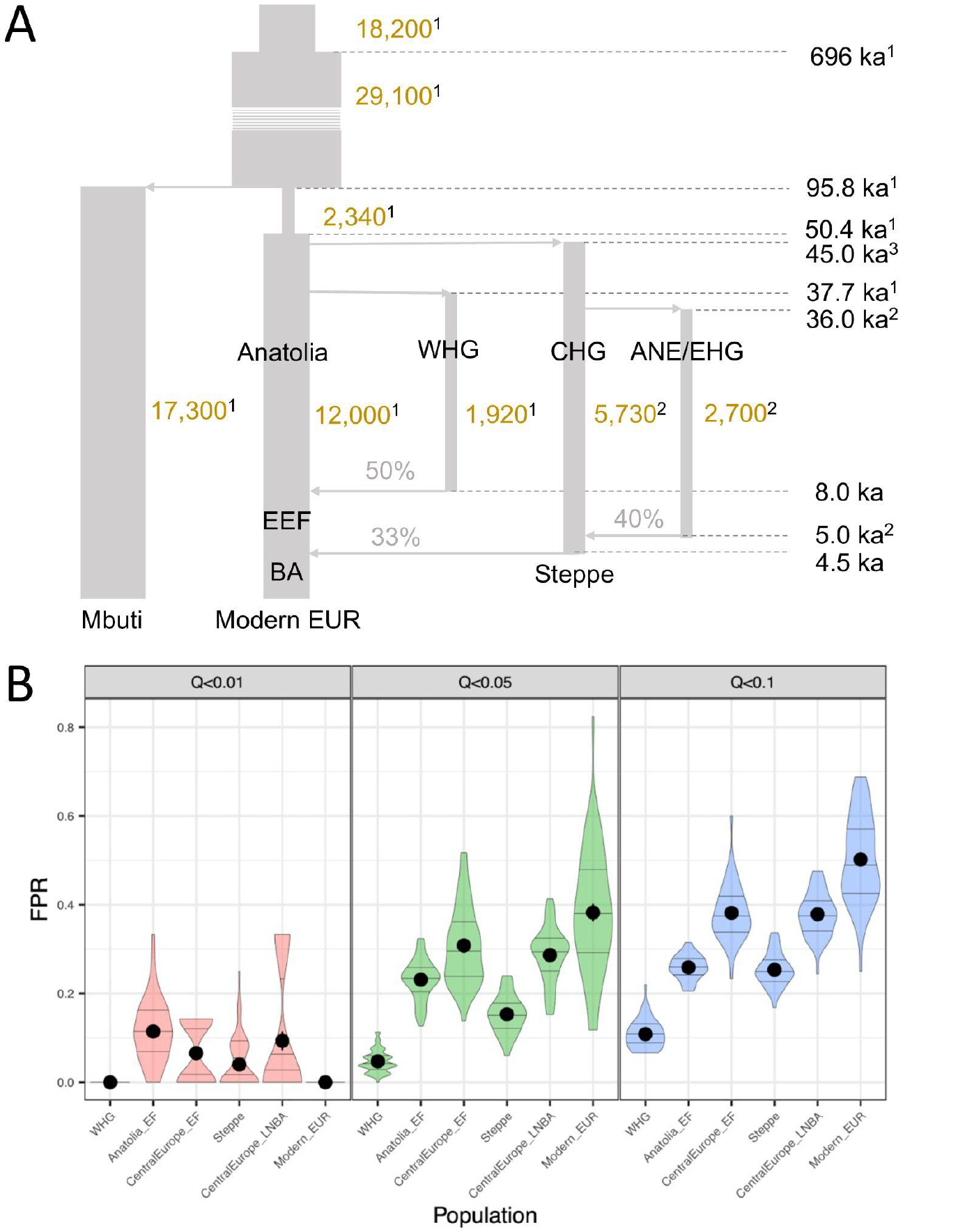
Assessing the robustness of the hard sweep detection pipeline. (A) Schematic of the West Eurasian population history model used to explore the statistical properties of our analytical pipeline and the impact of historical bottlenecks and admixture on the false positive rate. Each vertical segment denotes a major population branch (effective population sizes shown in gold text), with grey horizontal arrows denoting separation and admixture events (times shown on the right-hand side of the figure, assuming that admixture occurred 500 years after the onset of the migrations shown in Fig. 1; with percentages indicating the proportion of ancestry contributed by the incoming admixture branch). Model parameters are taken from one of three studies, as denoted by the associated superscript (1 = ref. ^84^; 2 = ref. ^33^; 3 = ref. ^34^). (B) Estimated false positive rate (FPR) measured at six different simulated populations sampled before (Anatolia_EF, Steppe, WHG) and after major admixture events (EF, LNBA, and Modern Europeans) using three different *q* value thresholds (see Fig. S19 for further information on sample size and sampling time). Results are shown for 30 simulated genomes, dots indicate mean values and horizontal lines represent quartile values. Candidate sweep detection was based on the most stringent threshold (*q* < 0.01; red densities) in the current study, whereby the maximum mean FPR observed amongst the simulated populations at this threshold (i.e. ∼11%) was used as a conservative estimate for the study-wide FPR.

### Hard sweeps were common in Paleolithic West Eurasian populations

In direct contrast to previous studies of modern human genomes^20^ we were able to identify large numbers of hard sweeps in the ancient West Eurasian populations (Figs. S8, S9, Table S2, Supplementary Data 1-57), identifying 57 with high confidence (estimated study-wide false positive rate of <11% under a realistic simulated Eurasian demography; Fig. 2; see Methods, SI Methods). None of these sweep signals are evident in the African Yoruba population, and ∼90% of the sweeps show significantly inflated levels of genetic divergence relative to African populations using *F*_st_-based tests (outFLANK method^21^; see Methods and Fig. S10), consistent with the underlying selection pressures postdating the separation of the founding Eurasian group from African populations.

Although these 57 putatively selected loci likely swept to high frequencies in West Eurasian populations by the Early Holocene to Bronze Age era (i.e. ∼12-5ka), only two sweeps (1:35.4-36.5 and 6:29.5-32.8; Fig. S8 and S9, Table S2) are still identified as SF2 outliers in selection scans of modern European populations, with these regions having been reported in previous selection studies^8^. This dramatic reduction in hard sweep signals was not an artefact of differences in either data quality or quantity between modern and ancient populations (Figs. S11-S14; SI Methods). Nor could their absence be explained by the degradation of the sweep signals through random allele frequency changes (i.e. genetic drift) and new mutations, as hard sweep signals in Eurasian human genomes are expected to remain visible to our analytical pipeline for around 70 thousand years in the absence of other effects (Fig. S15; see SI Methods and also refs. ^18, 22^). Rather, the loss of the hard sweep signals is consistent with the reduction in sweep haplotype frequency below detection limits caused by the introduction of other orthologous haplotypes during the Holocene admixture phases. For the analytical pipeline used in this study, power drops below 50% if the sweep haplotype does not occur in at least 85% of the population at the time of sampling, and becomes effectively undetectable if less than half the population has the sweep haplotype (Fig. S15 and SI Methods, also see ref. ^23^).

### A historical hard sweep in the MHC-III region is now masked in modern Europeans

Perhaps the most striking example of a previously unreported hard sweep in this study is provided by a ∼1.5Mb region overlapping the major histocompatibility complex class III (MHC-III) region on chromosome 6 that shows depleted genetic variation in the Anatolian Early Farmer (Anatolian EF) population (Fig. 3). The MHC-III region includes numerous genes that encode antigen proteins involved in the immune response and has previously been identified as a recurrent target of balancing selection^24–26^ and also as a potential source of selection artefacts due to the high levels of local genetic variation hindering read alignment across this region. However, nearly all (98.6%) of SNPs in the MHC-III sweep region lie in regions of high mappability, which is consistent with levels observed in the other 56 sweeps (minimum = 96.7%; Table S3), indicating that poor read alignment was not an issue for our sweeps in general.

**Fig. 3.**
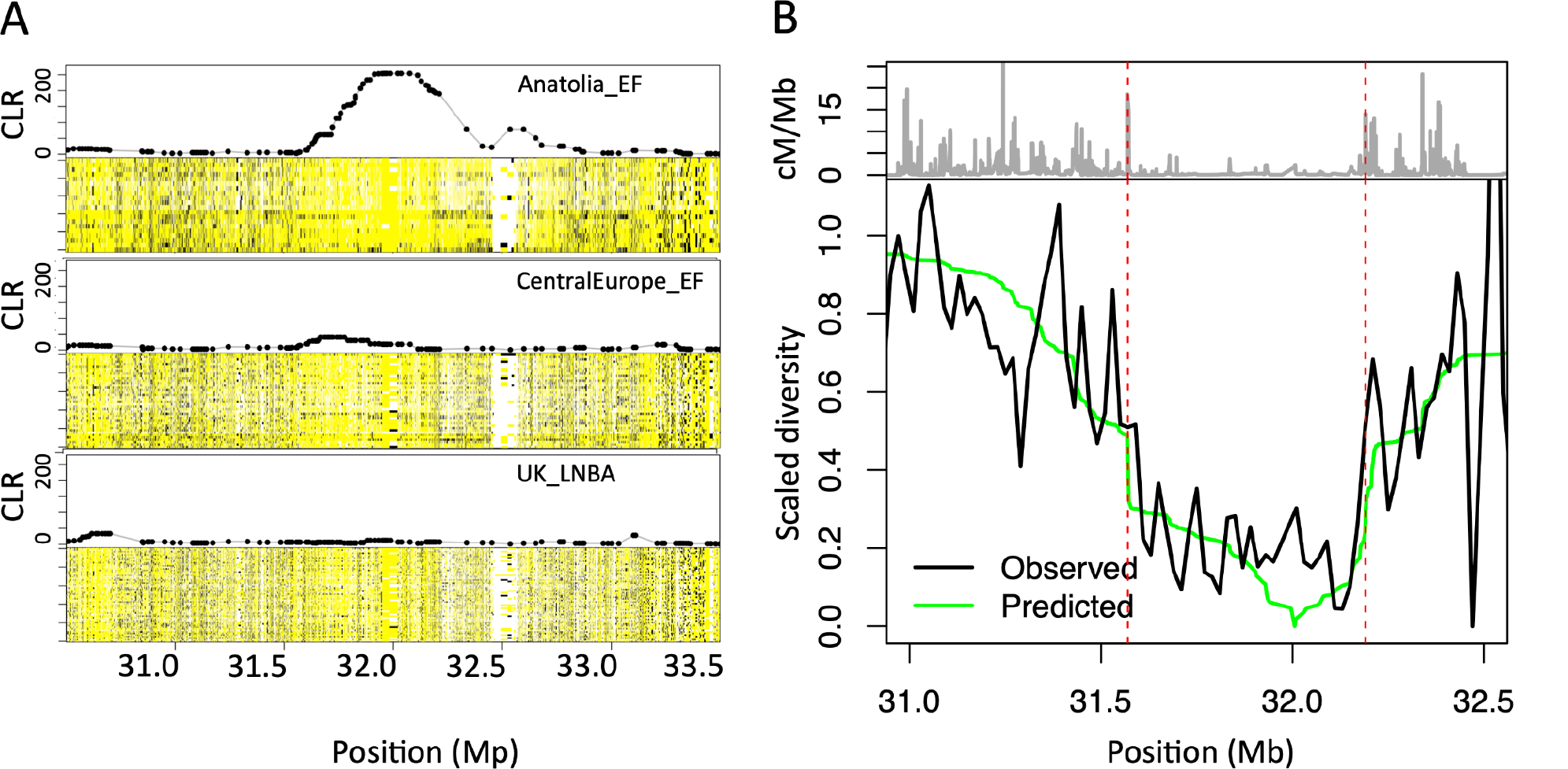
Hard selective sweep in MHC class-III region in Anatolian Early Farmers. (A) Haploimage of the MHC-III region and associated SweepFinder2 CLR score for the Anatolian Early Farmers, Central European Early Farmers (CentralEurope_EF) and British Bronze Age (UK_LNBA) populations. Pseudohaploid calls are shown for all samples in each population, with major alleles in yellow, minor alleles in black, and missing data in white. Elevated CLR scores coincide with a region of depleted variation in the Anatolian EF population, which returns to background levels in the subsequent admixed populations. (B) The estimated nucleotide diversity across the MHC-III region of the Anatolian Early Farmers (black line) and expected diversity under a hard sweep model (green line, see Text S3) relative to the underlying recombination rate in cM/Mb (grey line on top). The two dashed red lines indicate two local recombination hotspots flanking the sweep region. The close correspondence between the expected and observed genetic diversity across this region is unlikely to be a bioinformatic artefact and points to the authenticity of the signal.

The authenticity of the MHC-III hard sweep is further supported by the distinctive trough of genetic diversity observed across the affected region in the Anatolian EF samples, which is located between two recombination hotspots (i.e. areas of locally inflated recombination rates) that also demarcate the sharp transition to background diversity levels either side of the sweep region (Fig. 3). This correlation between genetic diversity and local recombination rates is unlikely to arise from read misalignment artefacts, but can be closely approximated by a hard sweep model acting upon a centrally located beneficial variant that incorporates the local recombination rates (Fig. 3; see Text S3), pointing to historical positive selection as the more likely cause.

In contrast to the Anatolian EF population, the two other major contributors to modern European genetic ancestry (i.e. WHG and Steppe) have SF2 scores and patterns of local genomic variation that are indistinguishable from neutral background levels in the same ∼1.5Mb MHC-III region (Fig. S16). Holocene-era populations show patterns that are consistent with a progressive dilution of the sweep signal across this period – with a weak SF2 signal still evident in Early European farmers (which draw ancestry from both Anatolian EF and WHG), that returns to neutral background levels in subsequent Bronze Age and modern European populations following the introduction of additional Steppe ancestry (Figs. 3, S16). Taken together, the evidence strongly implies that the MHC-III region was a target of strong positive selection in the Anatolian EF population, and had likely gone unnoticed previously due to the masking effects of Holocene-era admixture.

### Admixture can lead to misclassification of historical hard sweeps

The Anatolian EF-specific MHC-III hard sweep suggests that if Holocene admixture is responsible more generally for masking the 57 hard sweep signals in modern European genomes, then sweeps that were specific to only one of the admixing ancient Eurasian populations (e.g. where selection is driven by a population-specific, or local, pressure) should be particularly prone to SF2 signal dilution from Holocene admixture events. Conversely, sweeps occurring closer in time to the Out-of-Africa (OoA) migration are more likely to have signals that survive Holocene admixture events by virtue of being present at high frequencies in all admixing populations. To test if sweep signal presence was dependent on the sweep antiquity, we inferred the approximate onset of the underlying selection pressure by manually scanning five moderate-to high-coverage Upper Paleolithic individuals for evidence of the sweep haplotype – namely, Ust’Ishim ∼45ka^27^, Kostenki14 ∼37ka^28^, GoyetQ-116 ∼35ka^28^, Věstonice16 ∼30ka^28^, and El Mirón ∼18ka^28^ – assigning each sweep into one of five age classes based on the oldest sample in which the sweep haplotype is observed (Fig. 4; see Methods and Text S4). Concordant sweep age classifications were obtained using an alternate quantitative method (based on a set of diagnostic marker alleles; Fig. S17 and SI Methods), suggesting that our classifications should provide robust inferences for aggregated sets of sweeps from the same age class, even though individual sweep dates are likely to be less reliable.

**Fig. 4.**
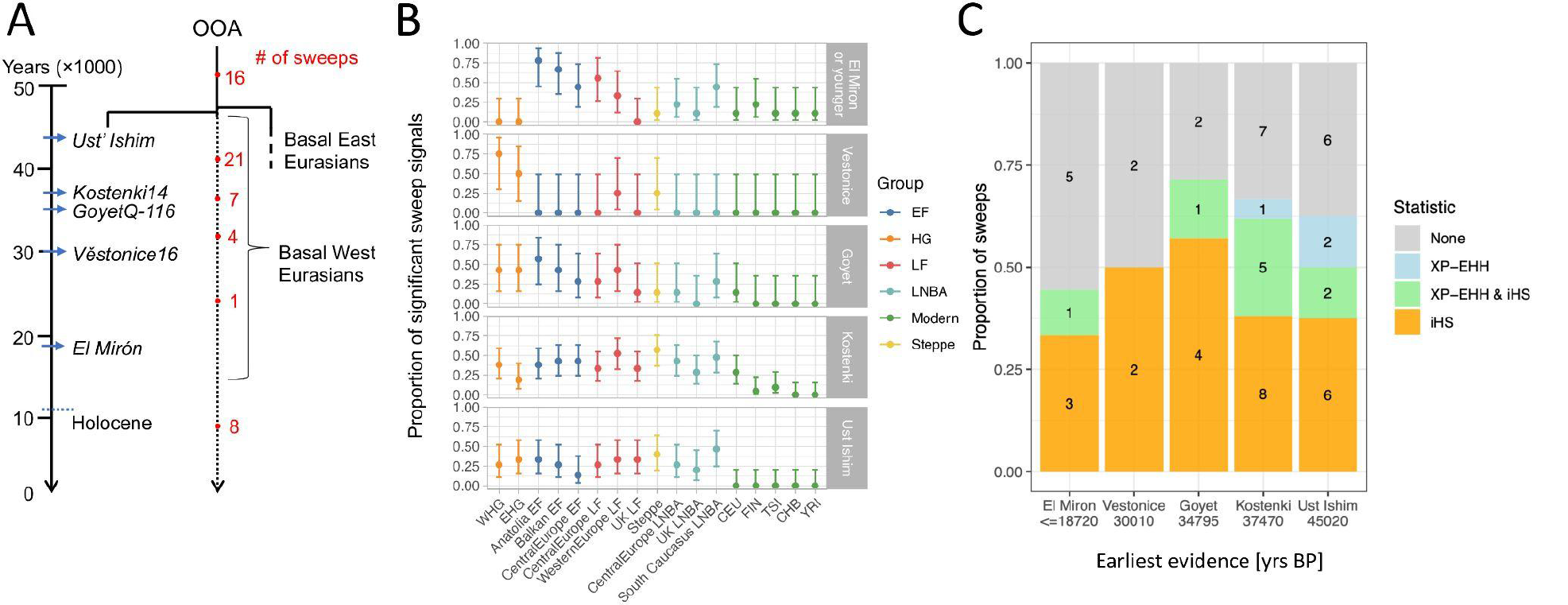
Older sweeps are more robust to population admixture. (A) Schematic representation of the inferred temporal origins of the 57 sweeps, classified according to the first presence of the sweep haplotype amongst the five moderate to high coverage Upper Paleolithic specimens (italicized labels, blue arrows indicating the approximate sample age). After classifying sweeps by their putative onset time, we calculated the proportion of sweeps present in each tested population (panel B; dots indicate point estimates at *q* < 0.05, error bars show 95% binomial confidence intervals) and in two studies reporting partial sweeps in modern Europeans^29, 30^ (panel C; iHS test statistics from ref. ^29^ being limited to outliers reported in at least two European populations to provide a stringent classification) conditional on the estimated onset times. Sweeps starting within the last 35kyrs (i.e. not observed in GoyetQ-116 or older samples) tend to have patterns consistent with local selection (i.e. being highly frequent in some ancient populations but absent in others; see Text S4) and are less likely to be reported as partial sweeps nearing fixation (i.e. exhibit an XP-EHH signal in ref. ^29, 30^; see key in panel C). While the latter difference was not significant (one-sided Fisher’s Exact Test *p* ∼0.17), our results are consistent with sweeps arising after the diversification of the Eurasian founders being more susceptible to admixture distortion

Sweep presence across the 12 ancient Eurasian populations was broadly consistent with the inferred antiquity of the selection pressure (Fig. 4B). Sweeps inferred to have started by 35ka (i.e. observed in GoyetQ-116 or an older sample) are detected at consistent levels across ancient populations both before and after the Holocene admixture events (i.e. no significant differences in sweep detection rates, logistic regression *p* > 0.17 for all three sweep age categories; see Methods and Text S4). In contrast, sweep detection differed significantly across the ancient populations for more recently selected loci (logistic regression *p* < 0.002 for both age categories) and exhibit patterns consistent with historical selection acting on populations ancestral to a subset of the tested populations (Fig. 4B). Similarly, sweeps starting by 35kya were more likely to appear as partial sweeps in two prior studies of positive selection in modern European populations^29, 30^ (66% vs. 46% for sweeps dated less than 35kya; Fig 4C), and were also more frequently observed as outliers for a statistic (i.e. XP-EHH) that is sensitive to loci nearing fixation (25% vs. 8%; Fig 4C). These results demonstrate that admixture can sufficiently distort the genetic signals resulting from a hard sweep, leading to the misclassification of the inferred mode of selection in studies where it is not explicitly accounted for.

### Evolutionary scenarios underlying sweep signal dilution

To further assess the plausibility of Holocene admixture events masking historical hard sweep signals in modern human genomes, we used population genetic simulations (SLiM3^31^) to model selection (Figs. S18-20; see Methods and SI Methods) within a plausible West Eurasian demographic model informed by ancient DNA studies^32–34^ (Figs. 2A, S19). Beneficial mutations were introduced on the Main Eurasian branch before (55ka) and after (44ka and 36ka) the diversification of the founding Eurasian population (Fig. 5, Figs. S19 and S20), with selection tests performed on the three ancient source populations (i.e. Anatolia EF and WHG both sampled at 8.5ka, and Steppe at 5ka) and three admixed European populations descending from two separate Holocene admixture phases^35^ at 8ka (i.e. European Early Farmers sampled at 7ka) and 4.5ka (i.e. Bronze Age Europeans sampled at 4ka, and Modern Europeans at 0ka; see Text S1). All simulated datasets matched the sample sizes, SNP numbers, missing data, and ascertainment bias observed in empirical data from the relevant populations (see Methods).

**Fig. 5.**
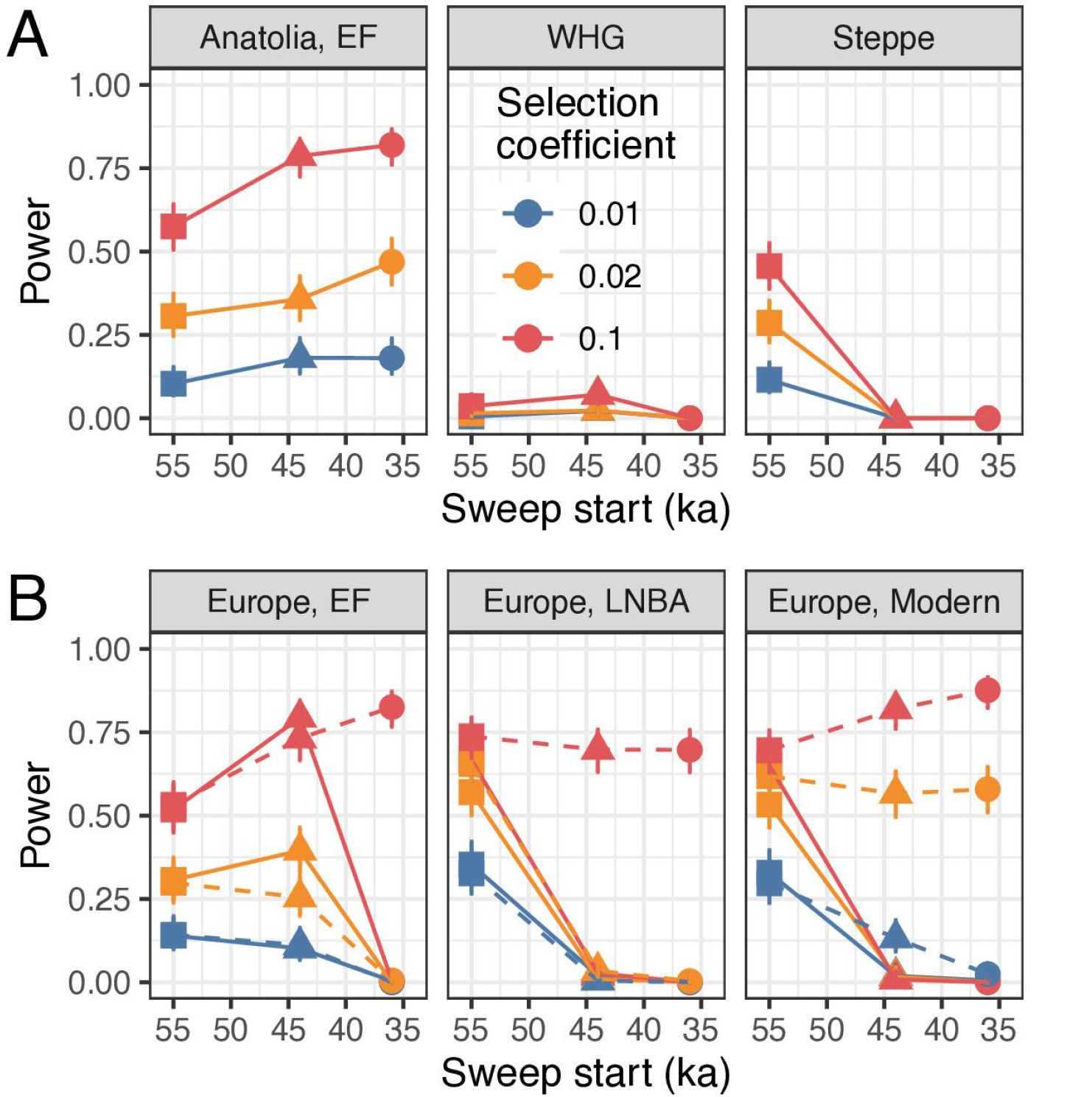
Investigating the influence of admixture on sweep detection in modern populations. Sweep detection power was estimated for selected loci simulated using a realistic Eurasian demographic model (see Figs. 2A, S19). Sweeps were timed to start prior to the diversification of the Eurasian founding population (55ka), or following the separation of population branches that eventually gave rise to Steppe (44ka) or WHG populations (36ka). Power was measured at a FPR of 0.1% and is shown relative to the onset of selection in (A) the three European source populations (i.e. Anatolia EF, WHG, and Steppe) as well as three admixed populations (B, C) following the mixing of WHG and Anatolian EF at 8.5ka (i.e. European EF, sampled 7ka) and European EF and Steppe herder admixture at 4.5kya (i.e. LNBA and Modern Europeans, sampled 4ka and 0ka, respectively). Only beneficial mutations preceding the diversification of Eurasian lineages (55ka) are evident in all populations when the selection pressure did not persist following the admixture events, with sweeps starting also 44ka also being detectable in European EFs since they are shared by both source populations (B, solid lines). Notably, power increases appreciably for strongly selected loci (*s* ≥ 0.02) when selection is allowed to continue in the post-admixture phase (B, dashed lines) due to these loci refixing following the admixture event.

We first investigated a model where selection is active along all population branches that inherit the beneficial mutation up until the 8ka admixture event, at which point the selection pressure is relaxed (e.g. due to an environmental change or the emergence/introduction of a technological innovation coincident with the admixture event). The results clearly demonstrate that Holocene-era admixture can effectively mask historical sweep signals in the absence of any ongoing selection pressures: only the sweeps that precede the division of Eurasian lineages (i.e. selection starting at 55ka) can be detected with reasonable power both before and after the admixture events. In contrast, sweeps starting after the diversification of Eurasian lineages (i.e. 44ka and 36ka) had slightly increased power in unadmixed populations that experienced the selection pressure (consistent with having more recent fixation times and less signal loss from drift), whereas power was negligible in all admixed European populations (Figs. 5, S20). As expected, the detection rates were strongly positively correlated with selection strength, whereas sample size had a more moderate positive association.

Interestingly, sweep detection power remained high in admixed populations whenever sweeps predated the split between the source population lineages – despite one of the source populations (WHG) having power close to zero under all modelled scenarios (Figs. 5, S20). The low power observed for WHG is likely a consequence of their relatively small effective population size^32^ exacerbating drift and causing rapid degradation of any fixed sweep signals (expected signal loss in 0.2 x *N_e_* generations^18^ = ∼11,000 years to loss in WHG assuming a generation time of 29 years^36^; *cf*. ∼70,000 years on the Main Eurasian branch). This result indicates that hard sweeps are reasonably robust to admixture-induced signal loss when the sweep signal has been eroded in one of the source populations due to drift, providing that the other source populations have retained the sweep signal.

After amending the model to allow the selection pressure to persist following the Holocene admixture events, we observe a dramatic increase in the detection rate of sweeps postdating the subdivision of Eurasian populations (i.e. selection onset at 44ka and 36ka), with power being particularly high for strongly selected loci (*s* = 10%) across all admixed European populations (power between 65-85%; Figs. 5, S20). Notably, Modern Europeans achieve detection power exceeding that observed for any of the three ancestral source populations (Anatolia_EF, Steppe and WHG) in nearly all cases. This result implies that many of the historical sweep signals should have been present in modern European populations had the underlying selection pressure persisted after the Holocene admixture phase, suggesting an attenuation of the underlying selection pressure(s), or the influence of other factors that were not included in the simulations, following the European Bronze Age (see Discussion).

### The mutational basis of the Eurasian hard sweeps

Under canonical models of hard sweeps, beneficial mutations only emerge following the environmental shift marking the onset of the selection pressure, though recent theoretical and simulation-based work has shown that hard sweep signals are possible when selection acts instead on mutations segregating at the time of the environmental shift (i.e. standing genetic variation; SGVs)^37–40^. Our results suggest that the underlying selection pressure(s) likely arose during the early stages of the AMH occupation of Eurasia – e.g. 16 [28%] and 44 [77%] of the 57 sweep haplotypes being observed in archaic samples dated at ∼45kya and ∼35kya, respectively (Fig. 4) – constraining the time in which *de novo* mutations could have arisen and suggesting that SGVs may have provided the mutational basis for many of our sweeps.

To examine this question in more detail, we calculated the probability of a sweep arising from SGV relative to a *de novo* mutation, conditional on the fixation of the beneficial allele within a particular time interval (see Text S5). Using equations from ref. ^41^ and assuming standard human generation times^36^, mutation rates^42^, and parameters from our demographic model (Table S4), we observe that fixation from SGVs is highly probable for selection strengths that match the majority of our empirical estimates (*s* ∼ 1%; see Table S2) when the temporal window in which fixation must occur is reasonably constrained (≤ 20kyrs), whereas *de novo* variants dominate when purifying selection on SGVs was strong (*s* ≥ 1%) in the period preceding the environmental shift (Figs. 6, S21). Extending our calculations to include the probability that selection from SGVs produces a hard sweep signal^41^ revealed that hard sweeps are highly probable (> 50%) regardless of the mutational input source, if SGVs had been deleterious in the period preceding the environmental change (Figs. S6, S21; Text S5). While further distinguishing between the two modes of selection is hampered by the absence of prior information on several key parameters (e.g. the strength of purifying selection on SGVs, the type and strength of epistasis, distance to the new fitness optimum), our results imply that mutations underlying the hard sweeps were probably initially deleterious and reinforce previous findings showing that selection from both *de novo* mutations and SGVs have occurred in Eurasian evolutionary history^39^.

**Fig. 6.**
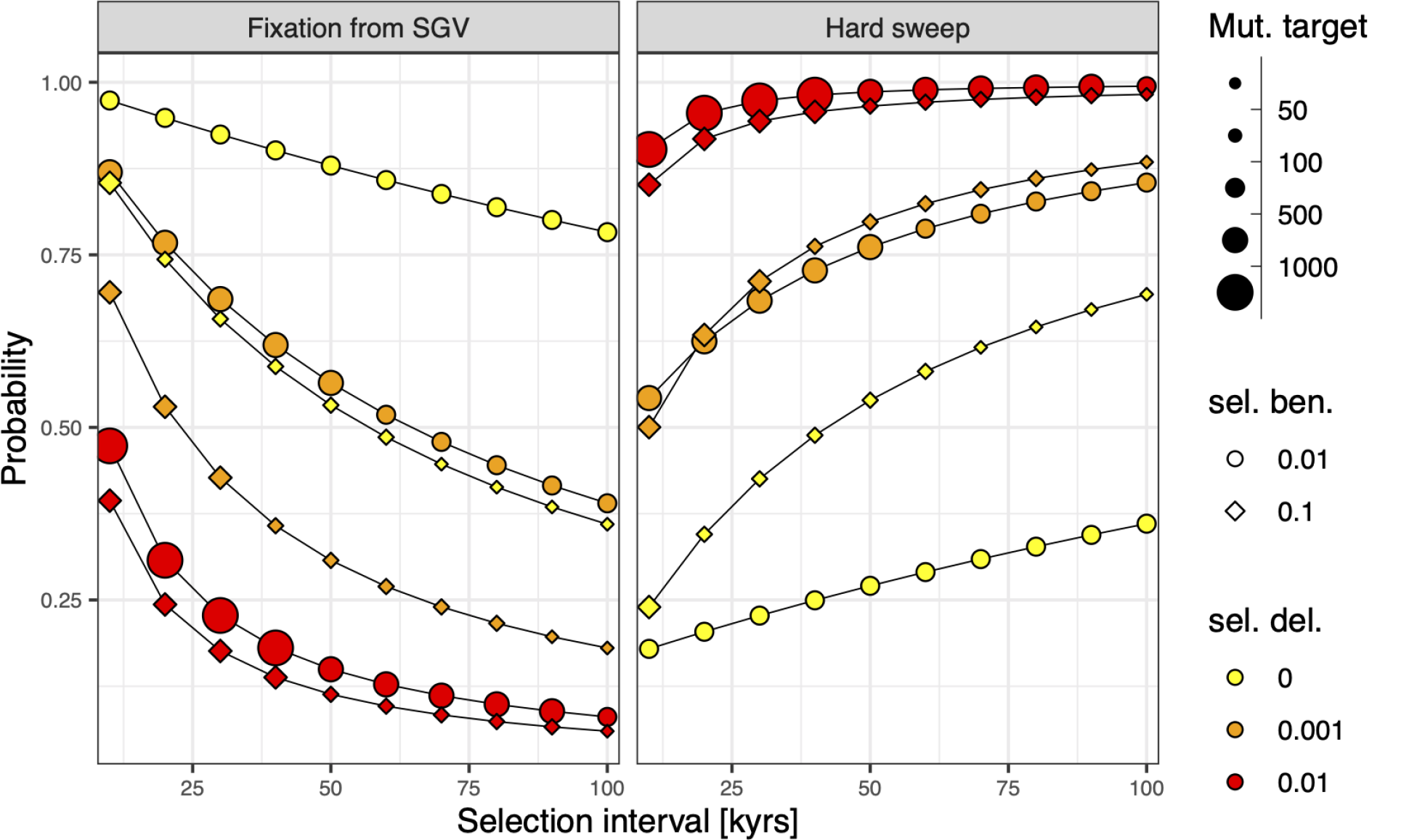
Mutational origins of the Eurasian hard sweeps. The probability that a sweep was caused by a standing genetic variant (SGV) relative to a *de novo* mutation conditional on a sweep of either type occurring within a fixed time interval (left panel), following equations in ref. ^41^. SGVs are assumed to have been under some degree of purifying selection (denoted by different coloured symbols) prior to the environmental shift that initiates the beneficial selection phase (symbol shapes). Fixation from SGVs was highly likely (>75%) for constrained time intervals (≤20kyrs) when the beneficial selection strength was moderate (*s* ∼ 0.01) – both being plausible values for Eurasian sweeps – provided that purifying selection prior to the environmental shift was weak (*s* ≤ 0.001). Factoring in the probability that fixed SGVs all descend from a single copy at the time of environmental shift results in consistently high probabilities (≥50%) that selection resulted in a hard sweep pattern regardless of the mutational source, provided that SGVs had previously been deleterious (right panel; assuming that all mutational targets arose in a single contiguous locus – noting that these probabilities will likely increase as mutational targets are distributed across more independent loci; see Text S5). Notably, the fixation of variants that had previously been strongly deleterious required large mutational target sizes (>1000 possible mutations) to ensure fixation within a 40,000 year interval when beneficial selection strength was ∼ 0.01, which may be implausibly large for many traits.

## Discussion

Our analyses of >1,000 ancient West Eurasian genomes has uncovered strong evidence for 57 hard sweeps in Early- to Middle-Holocene populations that have been almost entirely erased from descendent populations in modern Eurasia. Most of these selected loci had likely swept to high frequencies well before the Holocene era, which is supported by the strong selection strengths that were inferred for the sweeps (41 sweeps having *s* > 1%, with the largest value of *s* nearing 10%; Table S2; see Methods, Text S6). These selection coefficients are comparable with the strongest currently known for human populations (i.e. *LCT* locus having s between ∼2-6% in the recent history of European populations ^43, 44^), and suggest that such strong positive selection events have been much more common in recent human history than previously recognised. Our empirical and simulation results implicate Holocene-era admixture as the primary factor attenuating these historical sweep signals, which has led to these signals being missed or misinterpreted as other modes of selection in previous studies.

An intriguing implication arising from the simulations is that the selection pressure(s) underlying the sweeps may have eased during the Holocene period in some cases. This period marked the introduction of new technologies, diets, a stable warm climate, and living conditions that introduced novel selection pressures (i.e. selection on the *LCT* gene to reduce the costs associated with milk consumption in adulthood^44^) and may have also reduced the intensity of historical selection pressures underlying the 57 hard sweeps. Alternatively, these patterns could have resulted from the introduction of novel ancestry sources into Europe following the Bronze Age period^45–47^, which would have further diluted sweep signals and left insufficient time for their reappearance even if the selection pressure was still present. Indeed, this may explain why modern Tuscans and Finns had the fewest sweeps of any population in this study, with these populations descending from post-Bronze Age mixing events that involved African ^45, 46^ and Siberian^47^ groups, respectively (Fig. 4B).

In addition to masking historical sweeps in human populations, the obscuring effect of admixture might explain why species-wide selective sweep signals are rare in many species^7, 48^ while being abundant in others^49–51^. Crucially, recently admixed populations often lack detectable signs of structure, in which case admixture will not be correctly accounted for in any subsequent selection tests. For example, modern European populations have been considered sufficiently genetically homogeneous for the purpose of selection scans^29^, despite recent ancient DNA studies revealing that Europeans have multiple diverged ancestry components^52^. Indeed, modern genomic data is often insufficient to establish past admixture events, with widely-used PCA and ancestry decomposition methods being unable to detect historical admixture signals when suitable proxy populations for the admixing groups are lacking. Similarly, while admixture creates temporal variation in the coalescence rates of homologous genomic regions, these patterns can be equally well explained by historical population size changes in a single continuous population^53^. Accordingly, species with little apparent population structure may actually be more prone to the confounding effects of recent admixture than those exhibiting clear structure, and this may partly explain why species with distinctive population structuring often show strong local selection signals (e.g. Swedish *A. thaliana*^3^, African *D. melanogaster*^54^, *M. lychnidis-dioicae*^55^), whereas genetically homogeneous taxa or populations tend to lack fixed hard sweeps but harbour abundant partial or soft sweep signals (e.g. European humans^6, 29^, North American *D. melanogaster*^56^, *M. silenes-dioicae*^55^).

In accordance with previous work^4, 57^, our results emphasize the importance of incorporating historical population structure and admixture events into the null models of selection tests. If this information is not available – e.g. because DNA from ancestral source populations is lacking, the default scenario for most species – then the interpretation of historical selection signatures could be highly misleading and heavily weighted against the detection of historical hard sweep events. These factors imply that the extent of past hard sweep events have likely been underestimated in natural populations in general, biasing our understanding of the mode and tempo of adaptation in humans and other species.

## Methods

### Population designation

All ancient individuals used in this study were assigned to historical populations based on published analyses of their genetic relationships in combination with details regarding their archaeological context, with that temporal and spatial variability between individuals being minimised where possible (Figs. S1 and S2, Table S1). Our sample grouping resulted in 18 ancient populations occurring before and after the major Holocene admixture events that created the genetic landscape of modern Europe^16, 52^ (see Text S1). Additional testing using alternate sets of samples for two ancient populations showed that sweep signal detection was reasonably robust to changes in population sample configuration, with the close phylogenetic relationships across the samples providing a buffer to (see SI Methods; Figs. S13 and S14).

### Data collection and processing

To produce a robust dataset and avoid potential bioinformatic batch effects, the raw sequence read data for 1,162 ancient genomic datasets (Table S1) was retrieved from the European Nucleotide Archive and processed through the following standardized pipeline. To minimize the risk of modern contamination, the forward and reverse reads of the paired-end reads were merged (collapsed) using fastp^58^, and only merged reads were retained (modern data is more likely to comprise large DNA fragments that do not collapse). All collapsed reads were filtered for potential residual adaptor sequences and chimaeras using Poly-X with fastp^58^. The retained filtered set of sequence reads were aligned to the human reference genome (h37d) using the Burrows-Wheeler Aligner v0.7.15^59^. All mapped reads were sorted using SAMtools v1.3^60^ and then realigned around insertions and deletions and potential PCR duplicate reads were marked and removed using the Genome Analysis ToolKit (GATK) v3.5^61^.

Prior to variant calling, all remaining aligned reads were screened and base-calls recalibrated for aDNA postmortem damage using mapDamage2^62^. To further limit the impact of postmortem damage on variant calling^63^, bamUtil^64^ was used to trim 3 base-pairs from each of the 5’ and 3’ ends of each mapped read. From the resulting set of reads, pseudo-haploid variants were called at the set 1240k capture SNPs^65^ found on the 22 autosomes, using a combination of SAMtools mpileup^66^ and sequenceTools (https://github.com/stschiff/sequenceTools). Pseudo-haploidization of read data is a standard strategy in aDNA analyses, whereby a single read is randomly sampled at each prespecified SNP position^65^ in order to mitigate potential biases introduced by differences in coverage or post-mortem damage between samples^16^. The 1240k capture was developed to minimize ascertainment in non-African populations and was used to generate data for most samples used in the study, whereby concentrating on the 1240k variants ensured a common and robust set of variants for the subsequent analyses. The pseudo-haploid variant calls were converted from EIGENSOFT format^67, 68^ to binary Plink format using EIGENSOFT. Plink v1.9^69, 70^ was used to assign samples to the predefined populations (Table S1) and convert the variants to reference polarized VCF files, with correct polarization being checked using BCFtools^66^. Finally, a custom Python script was used to generate the site frequency spectrum (SFS) input files for SF2 analysis (https://gist.github.com/yassineS/fe2712ad52d76460b927e3f391ea51f6).

### Sweep scans

The SweepFinder2 composite likelihood ratio (CLR) statistic^18, 71^ was computed across successive 1kb intervals across all autosomes for each ancient and modern human population. The CLR statistic evaluates the evidence for hard selective sweeps in dynamically sized windows, by calculating the expected SFS under a hard selective sweep conditional on the neutral SFS, assuming a certain selection coefficient and local recombination rate. The neutral SFS is based on the ‘background’ SFS calculated from the whole genome assuming that the influence of positive selection on the genome-wide SFS is negligible.

SF2 controls for genome-wide effects such as ascertainment bias and demography by allowing these processes to affect the expected SFS under neutrality (i.e., the background SFS)^72^. Further, unlike many other selection methods, the assumptions on the input data for SF2 are suitable for the low coverage and ascertained nature of ancient DNA datasets. Since it is only based on the spatial (genomic) pattern of allele frequencies but not on haplotype homozygosity or population differentiation, it is possible to detect selection without reference to a second population, calling genotypes, or phasing haplotypes. By leveraging an empirical null model (i.e. the background SFS) and a model-based alternative hypothesis, SF2 is both more powerful and more robust than alternate test statistics also based on deviations of the SFS from expectations under the standard neutral model (e.g., Tajima’s *D*, Fay and Wu’s *H*)^18, 71^.

Note that SF2 also has an option to detect sweeps based on local genomic reductions in diversity. However, we did not calculate this diversity-based metric since the accurate and unbiased estimation of diversity requires fully sequenced genomes, whereas our dataset consists of an ascertained set of SNPs.

### Outlier gene detection

Human gene annotations were obtained from the ENSEMBL database^73^ (genome reference version GRCh37), which was accessed using the R biomaRt package^74, 75^ (version 2.36.1). Of the 24,554 annotated ‘genes’ on the biomaRt database, we removed any that were not annotated in the NCBI database (ftp://ftp.ncbi.nih.gov/gene/DATA/GENE_INFO/Mammalia/Homo_sapiens.gene_info.gz) and also excluded those that lacked specific protein and RNA based annotations (in the biomaRt transcript_biotype field). This resulted in a list of 19,603 genes, from which we removed 26 that did not contain any polymorphic sites in our datasets (all being situated in the most terminal areas of chromosomes), leaving 19,577 genes that were used in the subsequent analyses (Table S2).

To obtain *p* values for each gene, we transformed the raw SweepFinder2 CLR scores to *Z* scores using the following series of steps (Fig. S3). For each population, all SweepFinder2 CLR scores were logarithmically transformed and assigned to each of 19,577 genes by binning the transformed scores within the genomic boundaries of each gene. The gene boundaries were extended by 50kb on either side in order to also capture cis-regulatory regions. Because this typically resulted in several scores per gene, we took the maximum score to represent the evidence for a sweep involving that gene. Each gene score was corrected for gene length using a non-parametric standardization algorithm^76, 77^, resulting in the gene scores having an approximately standard normal distribution. Finally, *p* values were calculated for all genes and a *q* value correction^74^ applied for each population. The *q* value is a Bayesian posterior estimate of the *p* value that accounts for the expected inflation of false positives due to multiple testing^74^, whereby a *q* value of 0.01 implies a false discovery rate of 1% per population in this study.

### Candidate sweep classification

Sweeps were identified by determining a set of outlier genes across all populations, which were classified into sweep regions according to 1) the distance between the mid-point of neighbouring pairs of outlier genes (i.e. inter-gene distance) and 2) overlapping sweep regions between populations. Step 1 was performed independently for each population, whereby all outlier genes with midpoints that were less than a specific distance apart from the midpoint of a neighbouring outlier gene were collapsed into a single category. After generating the collapsed categories for each population, step 2 was applied to ensure that the sweep categories sharing at least one gene across different populations were considered as a single historical sweep.

We ran our sweep quantification pipeline at three *q* value thresholds (i.e. *q* < 0.01, 0.05 or 0.10; which imply false discovery rates of 1%, 5% and 10% per population, respectively) and the three different inter-gene distances (midpoint distances less than 250kb, 500kb or 1Mb). As expected, changing the *q* value had a large impact on the number of sweeps (ranging from ∼50 for *q* < 0.01 to ∼500 for *q* < 0.1), whereas changing the inter-gene midpoint distance had comparatively little impact overall, particularly at more stringent q-value cutoffs (Fig. S8). Based on these results, we decided to use the most stringent q-value cutoff and the most liberal inter-gene distance to define a conservative set of candidate sweeps for all further analysis. However, because this stringent cutoff might lead to the removal of potentially causal genes in a sweep (which could have values slightly lower than 0.01), we first defined our sweeps based on the more permissive *q* < 0.1 threshold, and then removed all sweeps that did not have at least one gene with *q* < 0.1. To further improve sweep determination, we removed populations with small sample sizes (see ‘Impact of small sample sizes on sweep detection’ below) from the sweep determination process, as previous analyses of modern genomes suggest that SF2 has little power to detect sweeps when the number of haploid genome copies being analyzed is 10 or less^23^. Finally, to ensure that the sweeps being defined were all relevant to western Eurasian history, the two modern populations from East Asia (CHB) and Africa (YRI) were also excluded from the sweep classification process. This strategy resulted in a total of 57 candidate sweeps that were used in subsequent analyses.

### Impact of small sample sizes and missing data on sweep detection

Previous results suggest that SF2 maintains high power to detect true sweeps when the number of haplotypes being analyzed is 10 or more for modern genomes^23^. To examine the impact of sample size on SF2 estimation on our ancient populations, we derived a measure of sample size that incorporates both pseudo-haploidy and different levels of data missingness in our ancient samples, called the ‘effective’ sample size, *n*_eff_. We calculated *n*_eff_ for each population as *kn*(1 - *M*), where *k* is the ploidy of each sample, *n* is the number of samples and *M* is the average proportion of missing sites at informative SNPs in that population. Consistent with results from modern data^23^, we found that the number of detected sweeps tended to be systematically lower for populations with *n*_eff_ values < 10, regardless of the *q* value threshold used (Figs. S8, S12).

When inspecting the distribution of *p* values for gene scores in each population, we observed large deviations from the idealized distribution as *n*_eff_ values decreased below 10 (Figs. S4, S5). These results indicate that populations with *n*_eff_ < 10 lack sufficient power to detect sweep signals, whereby only the 12 ancient populations with *n*_eff_ ≥ 10 were used to determine the candidate sweeps.

### Selection strength inference

We inferred the selection strength, *s*, for each of our 57 candidate sweeps as *s* = *r* ln(2*N*_e_)/α (from ref. ^78^), where *N*_e_ is the effective population size, *r* is the recombination rate, and α is a composite selection parameter that is estimated for each sweep region by SF2^18^ (see Table S2). For each candidate sweep, we took α at the position of the largest SF2 CLR value from all ancient Eurasian populations. Recombination rates were estimated for each sweep using information from ref. ^79^, and *N*_e_ was taken as 10,000 (noting that the estimation of *s* is robust to changes in *N*_e_ due to the logarithmic transformation). Notably, estimating *s* based on SF2 α estimates tended to result in slightly lower values than expected (see SI Methods and Fig. S18), suggesting that we have likely systematically underestimated the strength of selection for the 57 observed sweeps.

### Estimating the earliest evidence for selection

To estimate when the selection pressure(s) underlying each of the 57 sweeps may have first arisen, we manually inspected five Upper Paleolithic Eurasian human samples with moderate to high coverage genomes: Ust’-Ishim^80^, Kostenki 14^81^, GoyetQ-116^28^, Věstonice^28^, El Mirón^28^ for the evidence of the sweep haplotype. The oldest sample with evidence for the sweep is taken as the origin of the selection pressure, and therefore provides a coarse lower bound on the onset of selection pressure. The sweep was called as present only if the full set of alleles observed in the sweep region were observed in the sample, noting that genotype calls were only possible for Ust’-Ishim (coverage > 40x), with pseudo-haploid calls being used for the four other samples.

Because some sweeps contained complex signals with multiple peaks – which may represent the combination of two or more neighbouring selection events into a single sweep region (see Table S2 for a list of sweeps, and Supplementary Data Figures S1-S57 to show the distribution of SF2 CLR scores across all populations) – we concentrated on alleles underlying the strongest signal in each sweep. The plots used to discriminate the sweep haplotypes for all 57 candidate sweeps are provided in Supplementary Data Figures S58-S114 (https://adelaide.figshare.com/s/f00443ef535a8af2a61d). To test the robustness of these qualitative assessments, we compared our qualitative classifications with those obtained from a quantitative method. Details of this method and the comparison with qualitative results are provided in SI Methods.

### Testing sweep detection rate relative to inferred onset of selection

We predicted that hard sweep signals would be more prone to admixture-mediated loss if the onset of the selection pressure postdated the initial diversification of the Eurasian founding populations, as these sweeps are more likely to be ‘local’ to a subset of the source populations contributing to modern European ancestry. Accordingly, we reasoned that sweeps of deep antiquity should exhibit consistent detection frequencies across all ancient populations used in this study, whereas more recent sweeps should show inter-population variation that reflect their restriction to a subset of historical source populations. To examine this hypothesis, we quantified the proportion of sweeps present in each ancient population conditional on the inferred age of the sweep (see *Methods: Estimating the earliest evidence for selection*), then used the glm function in the R statistical language to test if proportions differed significantly across ancient populations grouped into five broad categories (HG, EF, Steppe, LF, LNBA; see Fig. 4). Specifically, for each sweep age class, we fit a logistic regression model where the presence of the sweep was a binary dependent variable and population group was the sole independent variable, and compared the fit of this model to the null model (dependent population variable removed, i.e. all groups have the same proportion of sweeps) to compute *p* values. Note that modern European populations were not included in these tests as our analyses indicate that these populations experienced further signal dilution following the Bronze age (see Text S1). Our results were unchanged after repeating all logistic regression analyses with the Southern Caucasus LNBA population removed from the LNBA group (since this population shows different ancestry composition to other European LNBA populations; see Supplementary Methods).

### F_st_-based selection tests

To further investigate the validity of the 57 candidate sweeps, we tested if our sweeps were enriched with highly divergent SNPs amongst the 12 ancient Eurasian populations with sufficient power to reliably detect sweeps (see Methods: Impact of small sample sizes and missing data on sweep detection) with a modern African population (i.e. YRI). We calculated *F*_st_ for each of the ∼1.1M ascertained SNPs using the standard Weir-Cockerham estimator^82^ and used OutFLANK^21^ to estimate the probability that each SNP was more divergent than expected under neutrality (based on estimating fitting a χ^2^ distribution to *F*_st_ values from putatively neutral SNPs). Importantly, OutFLANK is robust to non-equilibrium demographic models^21^, including rapid range expansions that are thought to have resulted in highly divergent alleles observed in modern human populations (i.e. through allele surfing^83^). Applying the *q* value correction to the *p* values to control for multiple testing resulted in 29 of the 57 candidate sweeps having one or more SNPs with *q* < 0.05, and 55 candidate sweeps having at least one SNP with *q* < 0.20 (Fig. S10). We then tested if the SNPs found in each of the 57 candidate sweeps had significantly higher *F*_st_ values than background genome levels. 49 of the 57 sweeps had significantly elevated *F*_st_ values relative to the remaining background genome (*p* < 0.05; Wilcoxon Rank Sum Test; Fig. S10), confirming that our sweeps were most likely selected following the divergence of ancestral Eurasians from African populations and were unlikely to be an artefact caused by non-equilibrium demographic processes such as allele surfing.

### Forward simulations

To assess the impact of Holocene admixture on historical hard sweep signals in modern human genomes, we used forward simulations (SLiM3^31^) to model moderate to strong selection (selection coefficient, *s*, between 1% and 10%) within a plausible West Eurasian demographic model (Fig. 2A and S19). Model parameters were largely taken from a recent study of Eurasian demographic history^84^, with further parameters for the Steppe population coming from two additional studies incorporating ancient Steppe samples^33, 34^. Two Holocene-era admixture events are included, namely a 50% contribution of WHG ancestry into the Main Eurasian branch at 8ka, and a further 33% contribution of Steppe (i.e. Yamnaya^33^) ancestry onto the same branch at 4.5ka. These admixture events generated the typical modern European individual that derives one-third of their ancestry from each of the Anatolian EF (modelled as descending from the Main Eurasian branch), WHG, and Steppe populations^16^.

Beneficial mutations were introduced on the Main Eurasian branch at three different times: 55 ka, 44 ka, and 36 ka (Fig. S19). We investigated a model where the strength of selection (i.e. *s* either 1%, 2% or 10%) was constant until the present, and another model where selection ceased following the 8ka admixture event (i.e. *s* = 0 after 8ka). Genomic regions of 5 Mb were simulated assuming mutation rates and recombination rates of 1x10^-8^/generation/bp, with the beneficial variant being introduced as a single mutation in the middle of the simulated region.

Diploid samples were generated for one modern population, sampled at the present, and five ancient populations: three populations that are ancestral to modern Europeans (i.e. Anatolia EF and WHG, sampled 8.5ka, and Steppe, sampled 5ka), and two admixed populations (i.e. Central Europe EF and Central Europe LNBA, sampled at 7ka and 4ka, respectively). Sample sizes matched those in empirical data, with 100 diploid samples taken for the modern Europeans. For each ancient population, we simulated missing data by randomly removing loci based on the site-specific missing data distribution of the relevant empirical samples, thereby replicating the population-specific data missingness patterns. We also reproduced the pseudo-haploidy of the ancient samples by randomly selecting one allele at each site. Additionally, because our analyses were limited to a set of ∼1.1 million variants from the 1240k capture probes (see Methods: Data collection and processing), which were selected based on heterozygosity in a set of sequenced reference individuals from both African and non-African populations^85^, we also sought to reproduce the ascertainment bias associated with these variants. To do this, we sampled two modern African samples and only retained sites that were heterozygous in either of the two African samples, or at least one of two randomly chosen modern European samples, with all other sites being discarded.

Finally, in all simulations, the number of sites was downsampled to 2000 positions in accordance with the average number of variants from 1240k capture probes found in a 5 Mb region. After removing all non-polymorphic sites, the SF2 statistic was computed at 5000 positions evenly distributed over the 5Mb region. To compute the neutral background SFS necessary for the SF2 analyses, we simulated 1000 neutral replicates of the demographic model and extracted the SFS for each population after replicating all steps in our simulation pipeline. Two hundred replicates were generated for each selection scenario and used for power analyses and to estimate the false positive rate (FPR). The final set of selection simulations are conditioned on sweeps that have escaped the initial stochastic phase (where rare beneficial alleles are lost through drift) by omitting all simulations where the beneficial mutation never reached a frequency of at least 10%.

More details on the statistical modelling procedure and full specification of the power analyses and FPR estimation are provided in the SI Methods.

## Author Contributions

A.C., Y.S., R.T. and C.D.H. conceived the study, Y.S., R.T., M.W. assembled the dataset, Y.S., R.T., A.J., C.D.H., G.G., M.C., A.R., and O.J. performed analyses, A.C. and C.D.H supervised analyses, Y.S., R.T., A.J., C.D.H., S.G., C.T., A.R., O.J., J.S., J.T., and A.C. interpreted the results, A.C., Y.S., R.T., A.J., and C.D.H. wrote the paper with input from all co-authors.

## Additional information

**Supplementary information** is available for this paper.

- Supplementary Methods
- Supplementary Text S1-S6
- Supplementary Tables S1-S4
- Supplementary Figures S1-S21
- Supplementary Data Figures 1-114

**Correspondence and requests for materials** should be addressed to Y.S., R.T., A.J., or C.D.H.

