## Supplementary Methods for "Admixture has obscured signals of historical hard sweeps in humans"

#### Population specification

To test the validity of population assignments, we performed a PCA analysis of the SNP data for each sample in the combined ancient and modern Eurasian populations. We used Plink v1.9[1] to prune the SNP set by removing closely linked variants (settings: `--indep-pairwise 500 kb 5 0.5`), and converted the resulting plink file to Eigenstrat *PACKEDANCESTRYMAP* format[2,3]. The PCA was performed using the *smartpca* function with the *noxdata, inbreed, numoutlier,* and *autoshrink* options all enabled, with the ancient samples being projected upon the PC axes defined by the three 1kGP European populations (*CEU*, *TSI*, and *FIN*). Examining the distribution of samples across the first two PC axes (Fig. S2) revealed that each of the ancient populations formed relatively distinct clusters and reproduced previous relationships reported for these samples (e.g. [4,5]), suggesting that they formed suitable approximations of populations for our analyses.

#### Outlier gene detection

For each population, all SweepFinder2 CLR scores were $log_{10}$ transformed and assigned to one of 19,577 genes by binning the transformed scores within the genomic boundaries of each gene (extended 50kb on either side in order to also capture *cis*-regulatory regions). As this typically resulted in several scores being assigned to each gene, we took the maximum score to represent the evidence for a sweep involving that gene. Each gene score was corrected for gene-length using the following non-parametric standardization function[6]:

$$Z_i = 0.675 \times \frac{y_i}{median(|y|)}$$

where $y_i = CLR_i$ - median($CLR$) and $i$ is an index for each gene in a given gene-length bin. The resulting set of adjusted gene scores approximated a standard Gaussian distribution – i.e. were $Z$ scores, with mean = 0 and standard deviation = 1 – whereby we used standardized Gaussian quantiles to compute $p$-values for all genes for each population (Fig. S3 and S4). Finally, to account for the expected inflation of false positives due to multiple testing, we applied $q$-value correction[7] to the *p*-values for each population. A *q*-value cutoff of 0.01 implies a false discovery rate of 1% per population.

Notably, when applying a one-sided test to determine outlier genes (i.e. those with scores lying in the upper tail), for a handful of populations the distribution of the resulting *p*-values was U-shaped rather than the J-shaped distribution that is assumed by the *q*-value correction method (i.e. a combination of non-selected genes that have a standard uniform distribution, plus a set of selected genes that mostly include low *p*-values). Hence, p-values were calculated by applying a two-tailed test to genes with $Z > 0$ (since selected genes will have larger positive $Z$ scores). This ensured that the expected *J*-shaped *p*-value distribution was obtained for each population (SI Appendix, Fig. S5). Importantly, this approach is not unnecessarily conservative, since our significance cutoff is based on *q*-values, i.e. the estimated false discovery rate (FDR) conditional on a specific *p*-value threshold. This ensures that our *q*-values will still be accurately estimated and correctly assign the FDR rate to the desired value.

**Coalescent simulation approach and issues**

Coalescent simulations provide an effective and time efficient way to simulate population genetic data, with extensions that allow modeling of selection at single locus (e.g., msms, [8]). Accordingly, we used the msms coalescent simulation software to simulate data under the demography reported in Fig. 6 (see section *Sweep detection power relative to start time and frequency at time of sampling* below). Counterintuitively, we found that our power to detect sweeps with SF2 increased after admixture from a population that had not experienced the selective sweep, relative to models where this admixture event was absent. Power is expected to decrease following admixture due to the dilution of the sweep haplotype frequency, which we subsequently confirmed when rerunning the demographic model using the forward-in-time simulator SLiM (Figs. 6, S16). Other msms users have also reported problematic results in demographic scenarios that combine selection and admixture (e.g. https://github.com/delt0r/msms/issues/41). Because msms is no longer being actively developed, we use the forward-in-time simulator SLiM[8] for all simulations in this study that include admixture (see Methods), using msms in a handful specific analyses where admixture was not modeled and the resulting simulated results were reliable (simulation type being noted in each relevant section).

**Modifications to the forward-in-time simulation framework to improve efficiency**

Of particular interest is measuring how the demographic history of West Eurasian populations impacts the false positive rate (FDR) and power of our analytical pipeline to detect hard sweeps. The demographic model and simulation framework is described in the Methods section. To speed up simulations, we generated 50 'burn-in' datasets, which comprised a single African ancestral population that was simulated for 220,697 generations, with a constant population size of 18,200 individuals from the start of the simulation about 6 million years ago until 696 ka, at which point the population size instantaneously increased to 29,100 individuals up until 95.8 ka (Fig. 6). These 'burn-in' datasets were randomly chosen to seed the subsequent simulated demographic history, substantially reducing the computational cost associated with investigating our highly parameterized demographic model with recent selective sweeps. To further expedite computation time, we rescaled the model parameters after the burn-in by a factor of four; specifically, we downscaled population sizes and time (in generations) by a factor of four, whereas mutation rate, recombination rate, and selection coefficients were increased by a factor of four. Because the beneficial mutation was often lost shortly after its introduction due to drift, further efficiency improvements were obtained by saving the state of the simulation at the point at which the mutation was first introduced and restarting all simulations where the beneficial mutation was lost from this saved state.

**Analytical pipeline false positive rate estimation**

Our analytical pipeline involves a series of steps that may lead to an inflated false-positive rate (FPR) if some of our implicit underlying assumptions are violated. In particular, we have assumed that the gene scores used in this study will follow a standard Gaussian distribution. To test if our assumptions were met and investigate the statistical robustness of our pipeline, we created and analysed 30 artificial population genomic datasets for each simulated population and demographic scenario. To create an artificial genome, SF2 CLR scores were calculated for the $n$ 5Mb sequences generated in each of the 2000 simulations, where $n$ is the sample size for a specific population (see Methods). Next, we randomly linked 565 of these 5Mb sequences into a full genome (noting that an individual 'sequence' in this context is the set of CLR scores computed from a single simulated set of samples). We then applied our analytical pipeline to each artificial genomic dataset: that is, we arranged the ~19k annotated genes used in the empirical analyses according to their linear position within the simulated genomic dataset, created individual gene scores (i.e. a standardised maximum overlapping CLR score; see Methods), determined outlier genes based on a $q$-value < 0.10, and then merged all outlier genes into sweeps by combining outlier genes less than 1Mb apart. Final candidate sweeps were determined by retaining those sweeps where at least one gene had a $q$-value below a specific threshold (i.e. either 0.10, 0.05, or 0.01). This process was repeated 30 times for each population and the FPR calculated for each $q$-value threshold by dividing the number of sweeps detected in each simulated genomic dataset by the number of sweeps observed in the empirical data at the same $q$-value threshold.

For all six simulated populations (Anatolia EF, WHG, Steppe, Central Europe EF, Central Europe LNBA, and Modern Europeans; see Methods and Fig. S19), the distribution of $p$-values for simulated genes closely match the expectations of the standard Gaussian distribution, with only a slight inflation in low p-values in all populations other than WHG (Fig. S5-7). The largest mean FPR was ~11% across the six simulated populations using the same q value threshold as our empirical analyses (i.e. $q < 0.01$), which is taken as a conservative estimate of the study-wide FPR (Fig. 2B).

**Testing the impact of recombination rate on sweep detection**

While SweepFinder2 is fairly robust to recombination rate variation [9], we performed separate msms coalescent simulations using the demography in Fig. S19, without the admixture events and omitting the CHG/ANE and EHG branches, under three different recombination rates: i) an 'average' rate of 1.45 cM/Mb (estimated by averaging across 1,000 randomly placed 1Mb windows using data from [10] and also ii) 2- and iii) 10-fold lower rates (0.72 cM/Mb and 0.145 cM/Mb, respectively). The two lowest rates were included to specifically test if the $p$ values associated with gene scores were systematically elevated in genomic regions with relatively low recombination rates, which could lead to an inflation of false positives at these regions. We performed 1000 neutral simulations for each recombination rate using the following msms command:

```
msms -ms 19 1 -oFP 0.00000000 -t 3000 -r $scaledRecRate -I 3 2
17 0 0 -n 1 0.95 -n 2 0.66 -n 3 0.11 -ej 0.018 3 2 -en 0.024 2
0.13 -ej 0.045 1 2 -en 0.045001 2 1.6 -en 0.330001 2 1
```

For each simulation, we generated multiple 5Mb sequences for four separate populations – Anatolia EF, WHG, UK LNBA, and Western Europe LNBA – replicating the missingness and sample sizes of each population (see more details in *Methods: Forward simulations*). We then generated 10 genomes for each population and used these to calculate the false positive rate following the same steps outlined in *SI Appendix Methods: Analytical pipeline power estimation*. The resulting distribution of *p*-values for simulated genes show a reasonable match to the expectations of the standard Gaussian distribution for all three recombination rates, with no systematic inflation of low *p*-values for lower recombination rates across the four tested populations (Fig. S7). These results indicate that genomic regions with low recombination were not more prone to false positives in our study.

**Analytical pipeline power estimation**

To test the power of our approach to detect selective sweeps in European history, we simulated positive selection under three different selection coefficients (*s* = 1%, 2%, and 10%) that covered the estimated range for our candidate sweeps, and three distinct onset times (55, 44, 36 ka; i.e. ranging from the early phases of the putative movement of AMH into Eurasia to shortly after the estimated split between the Main Eurasian lineage and WHG populations ~38ka [11] (Fig. S19). Two selection scenarios were modelled – both had a uniform selection pressure acting on all descendant population branches, with the first model assuming that selection acted until the present and the selection ceasing after the WHG-Main Eurasian admixture event at 8 ka in the second model.

Two hundred simulations were performed under each combination of the selection parameters (18 separate simulations in total; see Figure S20), with samples being drawn for six populations (i.e. Anatolia EF, WHG, Steppe, Central Europe EF, Central Europe LNBA, and Modern Europeans; see Methods). To create selected genomes, we first created a 'neutral' genomic scaffold by following the steps outlined in *SI Appendix Methods: Analytical pipeline false positive rate estimation*. After all genes were annotated and a score allocated to each following our analytical pipeline, a single gene was randomly selected and its score replaced with the maximum CLR score from one of the selected 5Mb segments. Standardized gene scores were then calculated using the neutral gene score distribution. This was repeated for each of the 200 simulations, with one of 50 neutral genomic scaffolds being randomly sampled to serve as the genomic background. Finally, we determined the upper 1% of gene scores for each neutral genome, and calculated the proportion of simulated gene scores that exceeded this threshold (comparing selected genes to the matching neutral genomic background). Performing this process resulted in 200 estimates of the power of our analytical pipeline (assuming a 1% FPR) for each of the six populations under each unique combination of tested selection parameters (Fig. 5 and Fig. S20).

Notably, the simulated WHG population displayed drastically lower power to detect sweeps than other Eurasian populations – being 20% units lower on average in the WHG population compared to the two 'large' ancient farming populations (Figs. 5, S20). Demographic parameters inferred by Kamm et al. used in our simulations suggest that the WHG population experienced a prolonged reduction in effective population size ($N_e$) compared to populations on the Main Eurasian branch (Figs. 2A, S19). The subsequent inflation in genetic drift caused by the lower $N_e$ in WHG lineages appears to have led to more rapid erasure of fixed sweep signals and a subsequent reduction of detection power relative to Main Eurasian populations.

**Testing the accuracy of SF2-based estimates of the selection coefficient**

We used msms to simulate selection along the Main Eurasian branch of the demographic model described in *SI Appendix Methods: Sweep detection power relative to start time and frequency at time of sampling*, ignoring all admixture events. Selection was simulated under four selection coefficients (1%, 5%, 10%, 20%) and two different starting times (33.8ka, 63.3ka). We further contrasted a model of selection from a new beneficial mutation (start frequency 1/2*N*) with a model of selection from a rare standing genetic variant (start frequency 0.1%). Following the same protocols as in our forward simulations, we simulated a 5Mb large region with the beneficial mutation occurring in the center of the region and replicated the sample size and missing data distribution of the respective empirical datasets (see Methods). The general msms command was:

```
msms -ms 19 1 -oFP 0.00000000 -t 3000 -r 2632 -I 3 2 17 0 0 -n 1
0.95 -n 2 0.66 -n 3 0.11 -ej 0.0142 3 2 -en 0.0202 2 0.13 -ej
0.0412 1 2 -en 0.041201 2 1.6 -en 0.326201 2 1 -N 18200 -SFC -SI
$start_time 3 0 $start_frequency 0 -SA $sel_coeff -Sp 0.5
```

We performed 200 simulations at each selection strength and starting time, with sweepFinder2 CLR scores calculated in successive 1000bp intervals along each 5Mb sequence and the selection coefficient calculated following the steps described in the *Methods: Selection strength inference* section. The resulting *s* values were compared to the simulated selection coefficients and are reported in Fig. S18.

Coalescent simulations to estimate power relative to start time and frequency at time of sampling

To determine how SF2 detection power varied with onset time of selection and the frequency of the sweep haplotype at the time of sampling, we used the coalescent simulator msms[8] to simulate 10,000 selected allele frequency trajectories along the Main Eurasian branch of our demographic model (Fig. 2A, S19), ignoring admixture events. Beneficial mutations arose at random within the past 200,000 years – initially along the branch that predates the separation of Eurasian and African populations, and subsequently along the Main Eurasian branch – with the population mutation rate assumed to be proportional to the current effective population size (Fig. 2A, S19). For each selected locus, we also simulated linked neutral genetic variation data across a 5Mb region for the hypothetical Early European Farmer population from 8ka (i.e. LBK branch in ref. [11]) and ran these sequences through SweepFinder2. To ensure that the simulated data match the statistical properties of the ancient data used in the present study, we replicated the empirical

SNP ascertainment scheme by conditioning on SNP presence in a simulated African population and also generated the same number of simulated 5Mb sequences as observed for the Neolithic Anatolian population in the present study (see Methods for more details on the ascertainment and missing data approaches in our simulations). Similarly, selection coefficients were sampled from an exponential distribution with a mean selection coefficient of $s = 1\%$, which leads to selection coefficients that broadly match those estimated from our sweeps ($s$ ranging between ~1 to ~10%; see Methods and SI Appendix, Table S2). For the subsequent analysis, we removed simulated selection coefficients of less than 1%.

To measure detection power based on the starting time of the sweep, we computed the percentage of simulations that had a SweepFinder2 CLR value larger than a 0.1% empirical FPR significance cutoff, conditioned on trajectories where the beneficial allele had reached a frequency ≥80%. The FPR was calculated as a function of selection starting time by using the default general additive model (GAM) in the R package *ggplot2*[12] smoothing function geom_smooth to calculate the best fit of a binary response (i.e. sweep CLR signals either significant or not) relative to selection start times for each range of selection coefficients (Fig. S15A).

To determine the relationship between detection power and the beneficial allele frequency when simulated 5Mb sequences were sampled at 8ka, we considered all final allele frequencies but only focused on sweeps starting within the past 20ka with selection coefficients > 1% (Fig. S15B). Here, we were primarily interested in the detection power of SweepFinder2 for incomplete sweeps that had insufficient time to reach fixation. Power was again computed using the geom_smooth GAM function, in this case computing the optimal fit of a binary response variable (i.e. significant CLR statistic based on a 0.1% FDR) relative to the final frequency reached by beneficial allele at the time of sampling.

Impact of sample sizes on sweep detection in modern populations

Modern populations tended to have systematically fewer sweeps than ancient populations in this study, a result that did not change when altering the $q$-value threshold used to determine outlier genes that define the sweeps (Figs. S8 and S11). To test if smaller sample sizes were more likely to produce more false positives (for example, by inflating the number of sweep haplotypes in a partial sweep by chance), we resampled one modern (FIN) and one ancient population with large sample numbers (Central Europe LNBA) to lower sample sizes (25, 10 and 5) and reran the complete detection pipeline to see if there was a systematic increase in the number of detected sweeps. We used an ancient population in addition to a modern population to take into account the potential impact of missing data and pseudo-haploidy on sweep detection. Notably, reduction in sample size did occasionally result in an increase in the number of detected sweeps for both ancient and modern populations; however, this effect was not systematic and in most cases the number of sweeps decreased in number – particularly at $n_{eff} < 10$ (Fig. S12). Further, the number of sweeps detected in the modern population was always lower than that detected for the ancient population at comparable effective sample sizes. Our results indicate that while there is some stochasticity in the number of reported sweeps, the overall pattern of decreased sweeps in modern populations is unlikely to be an artefact of larger sample sizes.

<u>Impact of population sample composition on sweep detection power</u>

To estimate the impact of our population assignment on the robustness of our candidate sweep detection, we reran the sweep detection pipeline on alternate sample groupings for both the *WHG* and *Steppe* populations and compared these to the original sample groupings. The original sample assignments for each population were governed by minimizing the temporal and spatial variability within each population while maintaining a homogeneous archaeological context. The alternate population groupings preserve the archaeological context but use a coarser geographical and temporal context, and/or include samples with additional ancestry components not present in the original sample groupings (Table S1). Specifically, for the *WHG* population – which initially comprised 44 samples largely derived from the Balkan region – we added 12 samples mostly sourced from the Gravettian culture from western Europe (sourced from France, Germany, and Luxembourg) that were ~1ka older on average (mean sample ages: 10ka vs 9ya for original samples). The 75 samples from the *Steppe* population were supplemented by 15 samples of similar antiquity and provenance (Maykop Culture samples; mean age ~4.5ka for both groupings), but which draw ~4% of their ancestry from Siberian hunter-gatherers (i.e. Eastern

Hunter-Gatherers with Siberian genetic affinity) that is absent in the samples in the original grouping[13,14].

The two sample groupings had significantly correlated gene scores for each population (Pearson's $r = 0.82$ and 0.64 for WHG and Steppe, respectively; Fig. S13). Notably, whilst changing the sample composition of the WHG and Steppe populations resulted in some different sweeps being detected for each (Figs. S13 and S14), 19 of the 27 sweeps (4/7 and 15/20 for WHG- and Steppe-specific sweeps, respectively) were observed regardless of which samples were used. Importantly, amongst these 19 retained sweeps, seven were observed in a population other than the alternate WHG or Steppe population (e.g. sweep *4:71.5-72.4* is significant in only one of the two Steppe sample groupings, however it is also significant in Anatolia_EF and CentralEurope LNBA; Fig. S14). Overall, 22 of the 57 candidate sweeps appear in two or more ancient Eurasian populations, which increases to 49 out of 57 candidate sweeps when sweeps are defined by containing at least one gene with a minimum $q < 0.05$ (instead of the $q < 0.01$). Our results imply that the shared genetic history and large number of ancient populations used in this study improve the chance that a sweep will be observed and thereby impart a degree of robustness to the sweep detection process.

Testing if sweeps were artefacts of low genome complexity

To test for potential false-positive sweep signals arising in genomic regions of low sequence complexity, we partitioned the genome into areas of high and low complexity based on CRG100 mappability scores[15] and checked if the proportion of mappable SNPs was systematically reduced in any of the 57 candidate sweep regions. Mappability is a metric that indicates how readily reads can be aligned to a particular genomic region, which has a positive association with sequence complexity. Hence, consistent observation of a low proportion of mappable nucleotides relative to the genome-wide average would suggest that many sweeps were false positives due to mapping artefacts. The CRG100 values were downloaded from the UCSC Genome Browser[16] and windows with a mean value less than 0.9 defined as a low complexity region, whilst remaining windows were defined as high complexity regions. For each of the 57 candidate sweeps, we calculated the proportion of ~1240k SNPs found within each sweep that also lie within high complexity regions. Notably, the SNP probes were designed to target genome regions with high mappability[17], and our results confirm this with at least 96.7% of the SNPs in each sweep lying in high mappability regions (Table S3). Taken together, our results imply that our candidate sweeps were not enriched with artefacts caused by low genome complexity, and likely represent true historical selection patterns.

**Sweep haplotype detection**

We qualitatively checked the occurrence of the sweep haplotypes in five Upper Paleolithic Eurasian human samples by comparing their haplotype plots with the dominant haplotype within the populations with significant sweep signals, for all 57 sweeps. To test the robustness of our qualitative method, we used the following quantitative method to infer the most likely selected haplotype. To reconstruct the sweep haplotype, we reasoned that the alleles carried on the selected haplotype would be at high frequencies in the ancient populations where the sweep was significant. Accordingly, for each sweep, we identified a subset of alleles that are strongly associated with the sweep, using the following two statistics:

$$\sum_{i}(1\text{-}4pq) \times ln(CLR_{\mathrm{i}}) \qquad (1)$$

$$ln[\sum_{i}p \times ln(CLR_{\mathrm{i}}) \,/\, \sum_{i}q \times ln(CLR_{\mathrm{i}})] \qquad (2)$$

Here, $p$ is the sample allele frequency, and $q$ is (1-$p$), for a specific SNP in population $i$. For each sweep, equation (1) was evaluated for every SNP lying in the sweep region to identify a subset of SNPs that act as reliable markers for the sweep. Equation (1) provides a weighted score for each SNP, which is the product of the log-transformed SweepFinder2 CLR score (i.e. the CLR score overlapping the SNP in question) and the standardised allele frequency, which is scaled to fall between 0 (when both alleles are at 50%) and 1 (when either allele is fixed). This value is calculated for each of the 18 ancient populations used in the study (indexed by $i$) and then summed together. Accordingly, equation (1) returns large weighted scores for SNPs where one allele is consistently near fixation in populations where the sweep also exhibits evidence for selection.

Once a subset of marker SNPs were identified for each sweep, equation (2) was used to evaluate the allele that is most strongly associated with the selected haplotype. Following the logic of equation (1), we calculated the weighted score separately for each allele in a given SNP, where $p$ and $q$ denote the frequency of each allele in population $i$, then calculated the log-transformed ratio of these two values. The sign of the resulting value indicates which of the two alleles is most likely linked with the haplotype (positive = $p$; negative = $q$), with larger magnitudes providing stronger evidence for association.

To investigate how the number and proportion of marker SNPs impacted our sweep classifications, sweep haplotype detection was based on the sample having at least 90% or 95% of the corresponding marker alleles, with sets of 50 or 100 marker SNPs being used. For cases where a sweep had fewer SNPs than the number of marker SNPs used in the analysis, all of the SNPs found in the sweep were used. Notably, one of the sweeps (*10:122.9-123*) only contained six SNPs overall (the next lowest had 23 SNPs in total), whereby the quantitative SNP haplotype inference was unreliable for this sweep and it was not used in the comparative analyses.

The correspondence between the quantitative results with the qualitative estimates for detecting the sweep haplotype in the five Upper Paleolithic samples for each sweep was evaluated by computing Pearson's correlation coefficient, $r$, for the complete set of sweeps (Fig. S17). Separate tests were run for all four combinations of the marker SNP criteria (i.e. haplotype presence based on either 90% or 95% of the top 50 or 100 marker SNPs). Additionally, we recalculated $r$ after removing 12 sweeps that had complex SF2 CLR signals that may have concatenated two or more sweep regions (Table S2 and Supplementary Data Figures S1-S57), since the quantitative method does not disambiguate these separate signals. We found good concordance between the qualitative and the quantitative method, particularly after removing complex SF2 CLR signals ($r = 0.43$ when based on 90% of 100 marker SNPs; Fig. S17), suggesting that sweep classification was reasonably robust.

### Supplementary Text

#### Text S1: Admixture in Holocene Europe

Previous population analyses of ancient genomes have revealed evidence of a complex history of Late Pleistocene European hunter-gatherer populations[18], which by the Early Holocene (~10-12ka) were distributed in a cline from the west (termed Western Hunter-Gatherers, WHG) to the east (Eastern Hunter-Gatherers, EHG)[19]. Around 8.5ka, a large migration of farming populations from the Anatolian region (Anatolian Early Farmers; Anatolian_EF) into Europe initiated a major period of genetic admixture with the extant WHG[19], resulting in Early European Farmer populations (Fig. 1). A second phase of admixture ensued over the next few thousand years, involving late surviving WHG groups and resident Early Farmers, and led to the emergence of Late European Farmer populations. The descendant populations subsequently underwent a further pronounced phase of genetic admixture with herders arriving from the Eurasian Steppe (Steppe) at the beginning of the Bronze Age, around 5ka[19], resulting in the farming populations of the Late Neolithic–Bronze Age period. Following the Bronze Age period, the emergence of successive large territorial empires encompassing much of Europe, the Near East, and North Africa (e.g. Neo-Assyria, Macedon, Roman, Ottoman) during classical antiquity (~3ka-1.5ka) initiated an unprecedented period of human movement[20,21] that facilitated further large-scale admixture across western Eurasia[22–26]. Later population movements, such as those associated with the Viking-Age expeditions starting around 1,300 years ago, further introduced and disseminated distinct ancestry sources around Europe[27], resulting in the complex genetic landscape that emerged during the European Current Era.

We note that the South Caucasus LNBA population contains substantial contributions from Caucasian Hunter Gatherer and Iran Neolithic populations that are much less frequent or absent in Holocene European groups[4,5,24,28,29]. Thus, while the South Caucasus LNBA are expected to be informative about Eurasian selection history, they are not representative of the admixture context specific to Holocene Europe and are treated accordingly in subsequent admixture-based analyses (with all exceptions being noted).

**Text S2: Sweep terminology**

The classical definition of a hard sweep is based on the fixation of *de novo* beneficial mutations and linked neutral variants[30]. However, hard sweep signals can still be created when the beneficial variant is rare, or the beneficial haplotype is near to fixation, so long as the underlying genealogy is sufficiently star-like[31]. In the present study, a hard sweep signal is created by a specific distortion of the site-frequency spectrum emblematic of a star-like genealogy (i.e. the elevation of rare and nearly fixed variants), which also encompasses selection from rare standing variation and partial sweeps near to fixation.

**Text S3: HLA locus sweep in Anatolian Neolithic samples**

The most striking SweepFinder2 signal in our data is a ~1.5 Mb wide sweep in the major histocompatibility (MHC) class III region of chromosome 6 in Anatolian Early Farmers. The MHC-III region is often overlooked in selection studies as it is regarded as a potential source of false positives arising from read misalignments caused by locally high levels of genetic variation. To examine the authenticity of the sweep signal further, we performed a series of additional tests examining if nucleotide diversity in this putative sweep matched expectations under a hard sweep model[32].

To estimate nucleotide diversity from our data, we calculated the expected heterozygosity for each SNP across the combined Anatolian Neolithic samples. Note that while our data is limited to a set of ~1.1 million SNPs and thus our diversity estimates do not estimate the true nucleotide diversity, these estimates can still be used to investigate local reductions in diversity as expected from a selective sweep. Accordingly, we calculated the average diversity in non-overlapping 20kb windows that span the sweep and flanking regions (from 23.5Mb to 35Mb on chromosome 6), rescaling diversity estimates by dividing them by the 90% diversity quantile across this region (since our raw diversity measure has no obvious interpretation).

We observe depressed levels of nucleotide diversity in a 1.5 Mb region centred directly beneath the SweepFinder2 signal peak, with diversity increasing sharply in the regions flanking each side of the sweep. Notably, this region of low nucleotide diversity also features relatively low recombination rates, with the transition to higher diversity being demarcated by two recombination hotspots (positions 31.57 Mb and 32.19 Mb; Fig. 4). This close correlation between regional recombination rate and diversity suggests that the depletion in diversity underlying the sweep signal is not a bioinformatic artifact related to read alignment problems or aDNA-related issues (e.g. data missingness), since these factors are unrelated to recombination rate and should not produce such a correlation. Rather, this relationship is consistent with the interplay between positive selection and recombination – i.e. neutral variants sitting near to the beneficial mutation remain in tight linkage while more distant variants lying beyond the two hotspots have become uncoupled by recombination during selection.

Notably, background selection could also lead to a reduction in local diversity in genomic regions marked by low recombination rate and a high density of functional elements. However, in this case we would expect that similar patterns would be observed in all populations examined in our study. Instead, most other populations exhibit weak SweepFinder2 signals and local allelic diversity consistent with neighbouring neutral regions (Figs. 3 and S16) – though the two European Early Farmer populations show pronounced (albeit insignificant) SF2 signals and some local diversity depletion that is consistent with having a high proportion of Anatolian Early Farmer ancestry along with some Western Hunter Gather ancestry.

Finally, we used the least squares method to fit a model predicting the reduction in diversity following the fixation of a selective sweep to the Anatolian Early Farmer samples. Following ref. [32], the predicted reduction in diversity relative to the genomic background was modelled as

$$\pi_{\mathrm{Sweep}}/\pi_{\mathrm{Background}} = 1- \exp[-r\log(2Ns)/s]$$

where $s$ is the selection coefficient, $r$ is the recombination distance between the neutral locus and the selected locus in Morgans, and $N$ is the haploid population size. We assume that the position of the beneficial mutation corresponds with the SweepFinder CLR maximum value (~32 Mb) and use a high-resolution human recombination map[33] to calculate the recombination distance of each SNP from this position. Fixing $N$ to 10,000, the best fit of this equation to the Anatolian Early Famer data was obtained when $s$ equalled 1.8%, with the predicted reduction in diversity for the Anatolian EF MHC-III sweep closely tracking the local nucleotide diversity patterns (green curve in Figure 3). Taken together, our results provide strong evidence that this region was a target of positive selection in the ancestors of Anatolian Early Farmers.

**Text S4: Inferred sweep haplotype presence in ancient and modern populations**

To test the role of admixture in masking historical sweeps in modern European genomes, we reasoned that the probability of a sweep signal to survive multiple bouts of Holocene admixture would be positively associated with the onset of the selection pressure underlying the sweep. Specifically, we manually checked five high coverage ancient West Eurasian samples for the presence of the beneficial haplotype for each sweep, and assumed that the timing of associated selection pressure was at least as old as the oldest sample exhibiting evidence of beneficial haplotype (see Methods and SI Methods).

After classifying each sweep according to these criteria, we found that around 77% of sweep haplotypes (44/57) are evident by ~35ka (i.e. the sweep haplotype was either present in Goyet, Kostenki, or Ust'Ishim), which we term 'Ancestral West Eurasian' (AWE) sweeps in recognition that the underlying selection pressures likely arose around the time of the OoA migration. In agreement with this proposed deep selection antiquity, the 44 AWE sweeps are detected at consistent levels (between 25-50%) across ancient Eurasian populations both before and after the Holocene admixture events. Intriguingly, the AWE sweeps show a marked decline in modern European populations (<20%; Fig. 5A), which suggests sweep signal dilution from admixture events that occurred after the Bronze Age (see Discussion).

In contrast with the AWE sweeps, the four sweeps that first appear at ~30ka (i.e. initially present in Vestonice) are common in WHG (~75%), but are less frequent in Steppe herders (~25%) and absent in Anatolia EF, and are also missing from all admixed ancient and modern European populations (Fig. 5A). These patterns suggest that these four sweeps most likely arose on the ancestral branch that links the Vestonice individual with WHG groups, following the separation of these lineages from the ancestors of Anatolian EF around 35-40ka[11], and thus are dubbed 'Local West Eurasian' (LWE) sweeps. These LWE sweeps are almost entirely absent from subsequent admixed European populations that draw their ancestry from both WHG and

Anatolian EF groups, supporting our hypothesis that local sweeps should be more susceptible to post-admixture signal degradation

Similarly, the eight sweeps in the final age class (≤19ka sweep class) have a pattern suggestive of local selection in Anatolia EF – i.e. 'Local Near Eastern' (LNE) sweeps' – being common in this population (mean ~75%) but rare in Steppe herders (~12%) and absent in WHG (Fig. 5A). LNE sweeps are also rare in Bronze Age and modern Europeans, pointing to post-Steppe admixture dilution – however, they are common in several Early and Late Farmer populations (~25-50%), a distinct difference from the LWE sweeps. This result suggests that incoming WHG ancestry contributions were insufficient to eradicate LNE sweep signals in some Early and Late Farmer populations. Alternatively, LNE sweeps may have had selection pressures that persisted during this admixture phase, and in one case (Sweep 5:144.9-145.9) the underlying selection pressure may have arisen during the Holocene (this being the only sweep not observed in any of the Anatolian EF, WHG, and Steppe populations at $q < 0.10$).

**Text S5: Hard sweeps from *de novo* mutations and standing genetic variants were both plausible in Eurasian history**

Our empirical and theoretical results suggest that hard sweeps were reasonably common events in West Eurasian history, echoing recent theoretical results that highlight how the demographic and evolutionary conditions inferred for founding OoA AMH populations were likely to have been conducive for the appearance of hard sweeps [34,35]. Specifically, one of the defining parameters for the appearance of hard sweeps are low to moderate values of the locus-specific population mutation parameter, $\Theta_l = 4N_e\mu$ (where $N_e$ is the effective population size and $\mu$ is the germline mutation rate), for selected loci in the period preceding the onset of selection. $\Theta_l$ has been estimated at $\sim 0{\cdot}001$ for non-African human populations [34,35], occupying the 'mutation poor' end of the parameter range where beneficial mutations tend to arise only after the onset of the selection pressure. Selection in this scenario tends to act on novel beneficial variants, which lead to long invariant DNA sequences that characterize hard sweeps following the fixation of the beneficial allele.

While novel beneficial mutations may have been the cause of several of the putative sweeps that we detect in ancient Eurasian populations, the fact that nearly 80% (44/57) of our hard sweeps were already evident in early Pleistocene samples originating prior to 35kya suggests that some of these sweeps may have arisen from standing genetic variants (SGVs) that were segregating at the time of the environmental change. Crucially, the waiting time for a new beneficial mutation, $t_w$, is inversely proportional to the census population size following the onset of the selection pressure[36], meaning that this was likely to have been prohibitively long in the current context unless the mutational target size underlying the selected trait(s) was very large.

To explore this question in more detail, we estimated the expected mutational target size necessary to ensure a sweep from either a *de novo* mutation or a SGV within a given time interval, $E[M|G]$ (i.e. the mean mutational target size, $M$, conditional on the number of generations, $G$, falling within a time interval in which the beneficial mutation must reach fixation, $t_f$), then used the estimated mutational target size to calculate the probability of fixation from SGV relative a *de novo* mutant, $P_{SGV}$. Specifically, following equations from ref. [37], we calculated the expected mutational target size, $E[M|G]$, as:

$$E[M|G] = \exp[-(\lambda_{SGV} + \lambda_{New})]$$

Where $\lambda_{SGV}$ and $\lambda_{New}$ measure the occurrence rate of mutants destined for selective fixation for SGVs and *de novo* mutants, respectively. These rates were obtained from equations 8 and 11 in ref. [37], resulting in the following:

$$E[M|G] = 1 / \exp[-(\Theta_d \log(1-R) + \Theta_b h_b \alpha_b g)]$$

where $\Theta$ is the population mutation parameter, $h$ is the dominance, and $R$ is the composite parameter, $2h_b\alpha_b / (2h_d\alpha_d + 1)$, $\alpha$ is the population selection parameter $2N_e s$ (with $s$ designating the selection coefficient), and $g$ is the number of generations rescaled in units of $2N_e$. The $d$ and $b$ indices denote whether the parameters occur before or after the environmental change that triggers the onset of selection, with $b$ indicating that the beneficial phase of the allele following the environmental change, and $d$ indicating the preceding deleterious phase (noting that the allele could also be strictly neutral during this phase, in which case $s_d = 0$). Importantly, this equation allows for different effective population sizes before and after the onset of the selection pressure, enabling bottleneck effects to be incorporated. We assume that the onset of selection coincided with the founding of the AMH migrant population after separating from ancestral African AMH, such that $N_{e(d)} > N_{e(b)}$.

After obtaining $E[M]$, we calculated $P_{SGV}$ following equation 14 in ref. [37]:

$$P_{SGV} = 1 - \exp[-\Theta_d \log(1-R)] / \{1 - \exp[-(\Theta_d \log(1-R) + \Theta_b h_b \alpha_b g)]\}$$

noting that Θ is now calculated as $4N_e\mu M$, i.e. the population mutation rate is rescaled according to the mutational target size calculated in the prior equation. All other parameters are recycled from the previous calculation.

Using these two equations, we estimated $E[M|G]$ and $P_{SGV}$ using standard estimates for μ (1.5 x $10^{-8}$)[38] and generation time (i.e. $g = 29$ years, noting that $G = t_f/g$)[39], setting dominance to be strictly additive (i.e. $h_d = h_b = 0.5$), and taking a range of values for other parameters that were not known *a priori* (see Table S4). The range of beneficial selection coefficients replicating estimates from the present study (see Methods and Table S2), with the ancestral $N_e$ set to 20,000 and Eurasian $N_e$ varying between 2,000 to 20,000 to model differential bottleneck effects, aligning with values in our demographic model (Fig. 2A and S19). Our results indicate that sweeps from SGVs and *de novo* mutations both were quite likely in Eurasian history (Figs. 6, S21), with *de novo* mutations dominating when purifying selection was strong ($s_d = 0.01$) prior to the environmental change, whereas SGVs were more likely when purifying selection was weaker ($s_d \leq 0.001$). Fixation from SGVs was particularly likely ($P_{SGV} > 0.75$) when the bottleneck intensity was strong (10% of ancestral $N_e$, i.e. 2,000), the fixation interval was constrained to ≤20kyrs, and selection strength during the beneficial phase was ~0.01, all of which are highly plausible for the observed Eurasian sweeps.

Next, we calculated the probability that selection from SGV resulted in the fixation of loci descending from a single variant at the time of the environmental shift, $P_{single}$, over the same set of parameters, obtaining an estimate of the proportion of fixed SGVs that guarantee a hard sweep signal. We calculate $P_{single}$ as 1 - $P_{mult}$, where $P_{mult}$ is the probability that multiple copies are fixed as defined by equation 18 in ref. [37]. As expected, $P_{single}$ exhibits a reverse dependency on the strength of purifying selection than $P_{SGV}$, directly reflecting the greater expected abundance of SGV copies available at the time of environmental change when purifying selection is weak (Figure S21). Nonetheless, the probabilities of a single copy fixing following the environmental change were non-negligible across the full parameter space explored, even when mutations were strictly neutral prior to the environmental change (minimum $P_{single}$ ~10%).

Finally, we calculated the compound function $P_{hard} = P_{SGV}P_{single} + 1 - P_{SGV} = 1 - P_{SGV}P_{mult}$, which estimates the probability that a fixed variant arising from either a SGV or a *de novo* mutation produces a hard sweep (i.e. the product $P_{SGV}P_{single}$ estimating the probability of a hard sweep from SGVs and 1 - $P_{SGV}$ estimating the probability of a hard sweep from *de novo* mutations, assuming that all fixed *de novo* variants produce a hard sweep). The inverse dependency between $P_{SGV}$ and $P_{single}$ on the strength of purifying selection meant that hard sweeps were highly likely when causal mutations experienced some degree of purifying selection prior to the environmental shift (minimum $P_{hard}$ ~50% when $s_d > 0$), with soft sweep signals becoming particularly dominant when causal mutations were previously strictly neutral and the fixation time intervals were tightly constrained (Fig. S21).

Notably, the $P_{single}$ calculation only considers scenarios where all beneficial mutations arise in a single locus, whereby we expect that the probability of a hard sweep signal from SGV could be higher if mutations occur across multiple independent loci. While a recent study suggests that selection on variants distributed across multiple loci tend to result in polygenic signals when $\Theta >$ 0.1, this work assumed that the fixation of a single beneficial mutation was sufficient to arrive at the new fitness optimum[35]. Accordingly, scenarios where the fitness optimum is more distant could support outcomes where one or more loci reach fixation when $\Theta$ exceeds 0.1. However, little *a priori* information is available to inform the parameterization of appropriate polygenic selection models for Eurasian history, including the distance to new fitness optima, mutational target size, epistasis between beneficial loci, and the strength of purifying selection prior to the environmental change, amongst others. Nonetheless, our results suggest that beneficial variants fixing in Eurasian history were likely to leave a hard sweep signal, lending further credibility to the 57 hard sweeps observed in this study.

**Text S6: Selection was sufficiently strong to result in several sweeps during the Eurasian Pleistocene**

Using the five Late Pleistocene individuals to time the candidate sweeps suggests that the majority of candidate sweep haplotypes were at high frequencies by 35ka, suggesting frequent episodes of surprisingly strong selection prior to the early Pleistocene period. Our estimates of $s$ indicate that selection was reasonably strong, with 53 sweeps having $s > 0.5\%$ and 41 having $s > 1\%$, with the largest value of $s$ nearing 10% (see Table S2). The selection coefficients we observe are strong enough to drive new or initially rare mutations to a detectable frequency (i.e. >90%) in only 3-4 thousand years for the strongest selection coefficients observed in our data, and within 10-20 thousand years for sweeps with the weakest selection coefficients (Fig. S18). Importantly, these estimates are likely lower than their true $s$ values – the SFS-based approach of SweepFinder2 tends to underestimate $s$ by about 20-30% [40], and selection from standing variation can lead to substantially narrower sweep regions, also resulting in underestimation of $s$. Indeed, when comparing the inferred selection coefficients against the simulated values for the *WHG* and *Anatolian_EF* populations, $s$ was consistently underestimated across the different selection regimes, particularly when acting upon more recent standing variation (SI Appendix, Fig. S18). Thus, our data are consistent with multiple episodes of moderate to strong selection driving beneficial variants to high frequency prior to 35kya, which were likely related to the new environmental conditions encountered by AMH following their arrival in Eurasia.

### Supplementary Figures

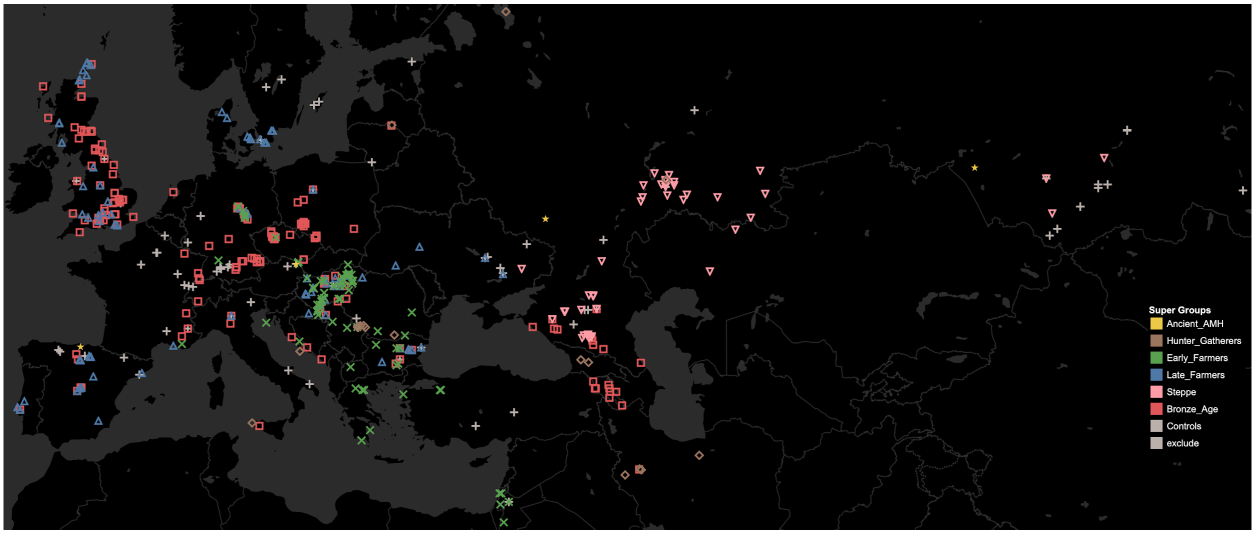


**Fig. S1.** Full geographic distribution of ancient Eurasian samples, including the far-Eastern Eurasian samples omitted from Fig. 1, with different populations (indicated by colored symbols) classified into broader groupings according to archaeological records of material culture and lifestyle.

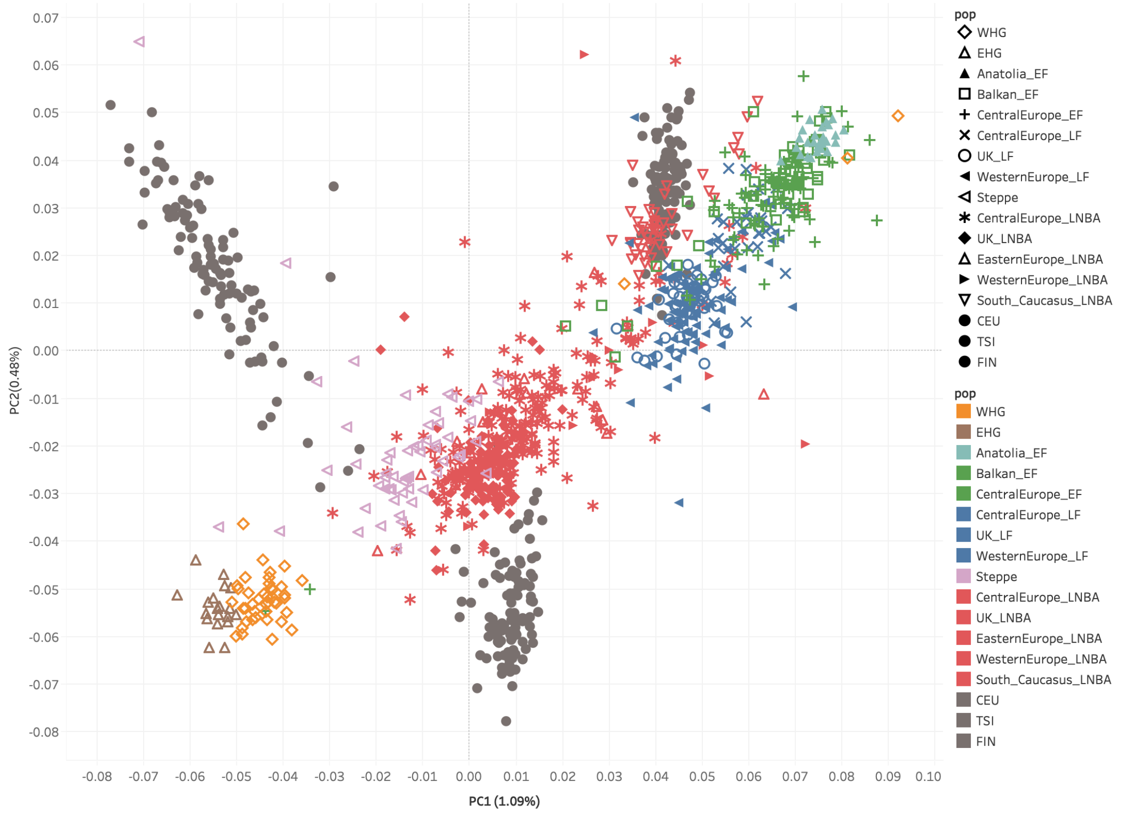


**Fig. S2.** PCA analysis showing the genetic relationships between the different populations used in this study with all ancient populations distinguished (see key). Ancient samples were projected onto modern samples from the three modern European populations (grey dots). See SI Methods for a full description of this analysis.

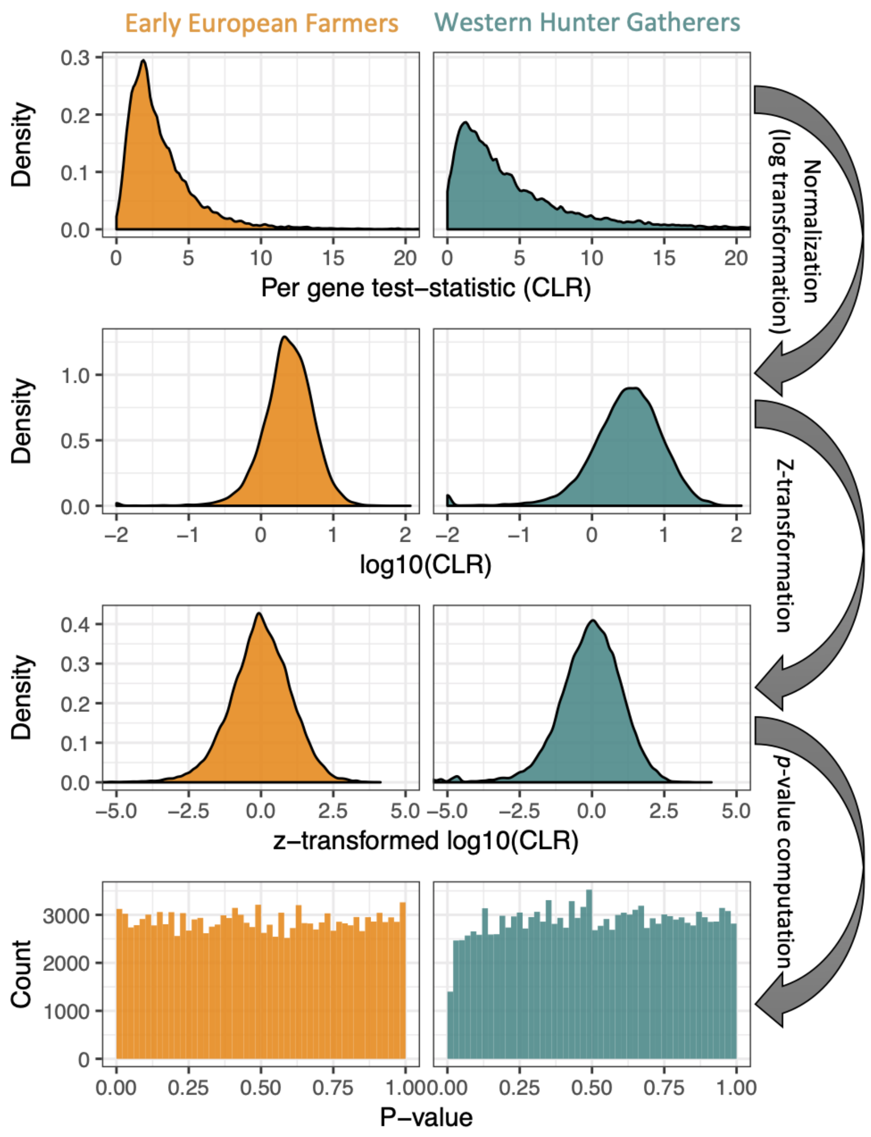


**Fig. S3.** Schematic representation of the series of transformations performed on the raw SweepFinder2 CLR test statistics to infer *p* values. The top panel shows CLR scores from two Eurasian populations simulated under a neutral demography (see Figs. 2A, S19), following the assignment of a single CLR score to each gene and subsequent correction to account for the positive correlation between gene size and CLR score (see Methods). Gene scores are initially approximately log-normally distributed, whereby subsequent logarithmic transformation produces normally distributed values (2nd panel) that are further centred and scaled to generate standardised (i.e. *Z*) scores (3rd panel), from which *p* values can be derived (bottom panel).

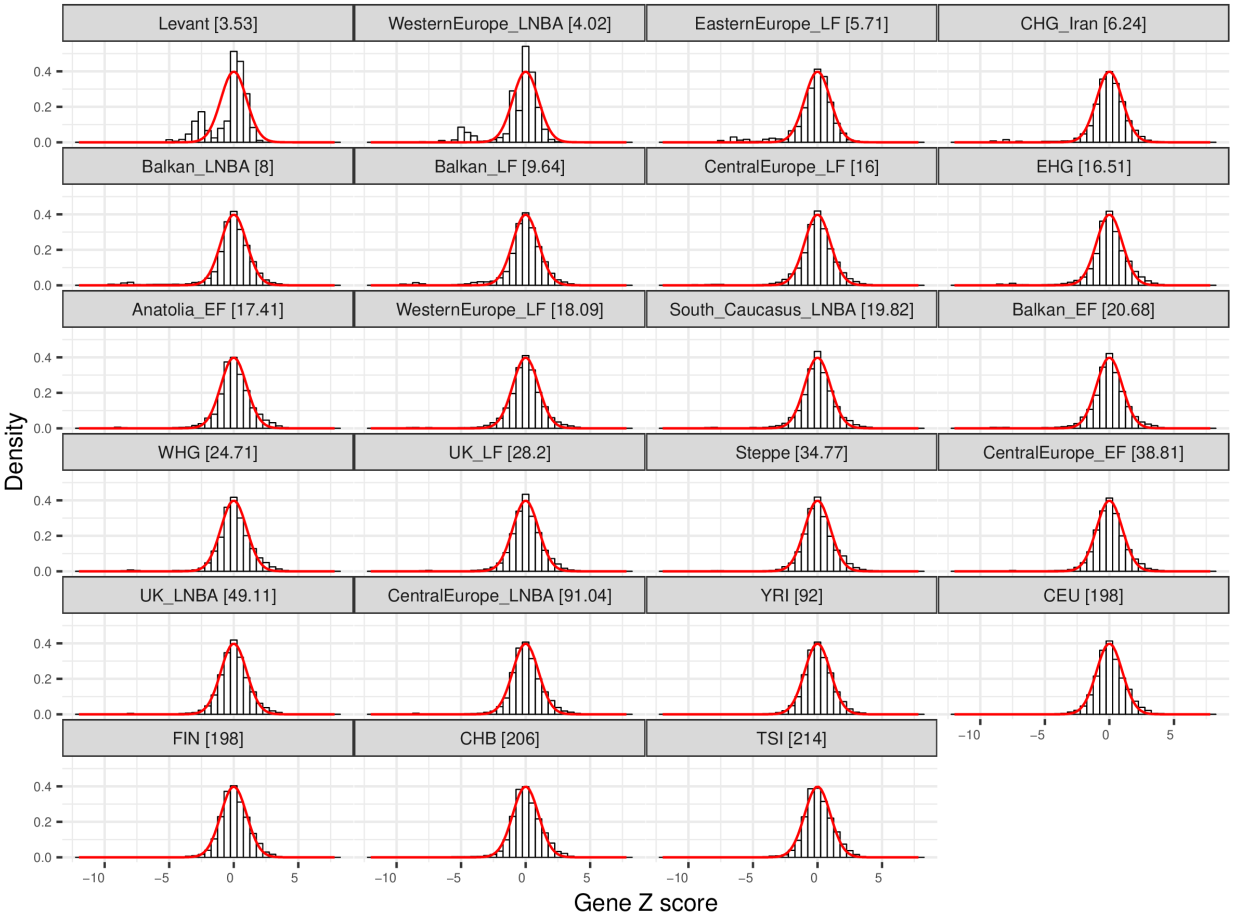


**Fig. S4.** Distribution of gene scores across all empirical populations. For each population, gene scores were generated following the process shown in Fig S3 (see Methods). All gene scores have an approximate standard normal distribution (*i.e.* Z scores, expected distribution = red line), though deviations become more evident as the effective sample size, $n_{eff}$ (a measure of sample size that accounts for pseudo-haploidy and missing data; see Methods), declines below 10 ($n_{eff}$ values shown in panel header in square brackets). Similar patterns were observed in the simulated datasets (Figs. S6 and S7), whereby only the 10 ancient Eurasian populations with $n_{eff} \geq 10$ (and the three modern European populations) were used to determine the 57 sweeps identified in this study.

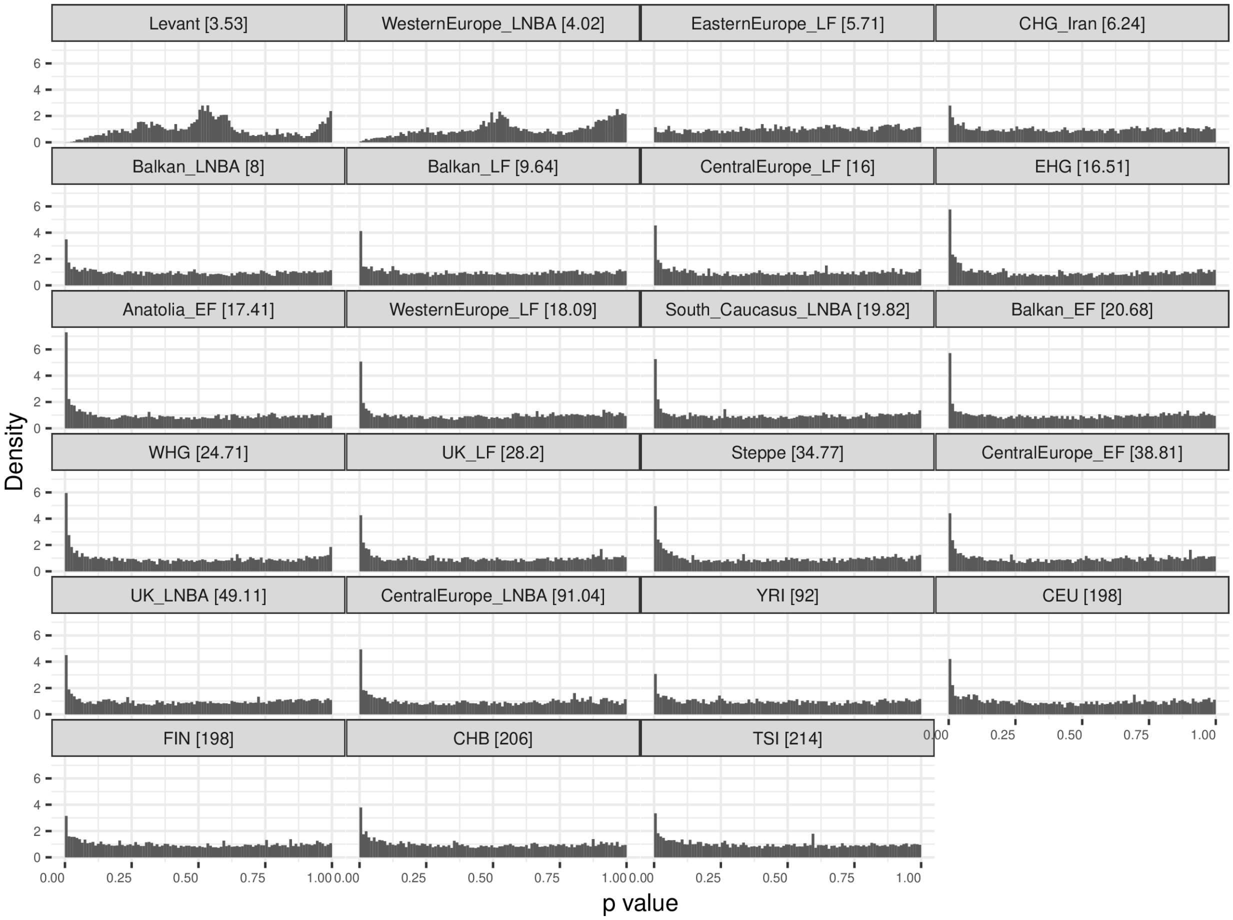


**Fig. S5.** Distribution of $p$-values for each gene, across all populations. Because some populations had U-shaped distributions, $p$-values were only calculated for genes with a $Z > 0$, noting that this did not impact outlier gene detection (see Methods). The J-shaped distribution observed for most populations implies that they contain a mixture of neutral genes (which have a standard uniform distribution) and selected genes (which cluster at low $p$-values). Deviations from the expected J-shaped distributions become more notable as the effective sample size, $n_{\text{eff}}$, declines below 10 ($n_{\text{eff}}$ shown in square brackets in the panel header).

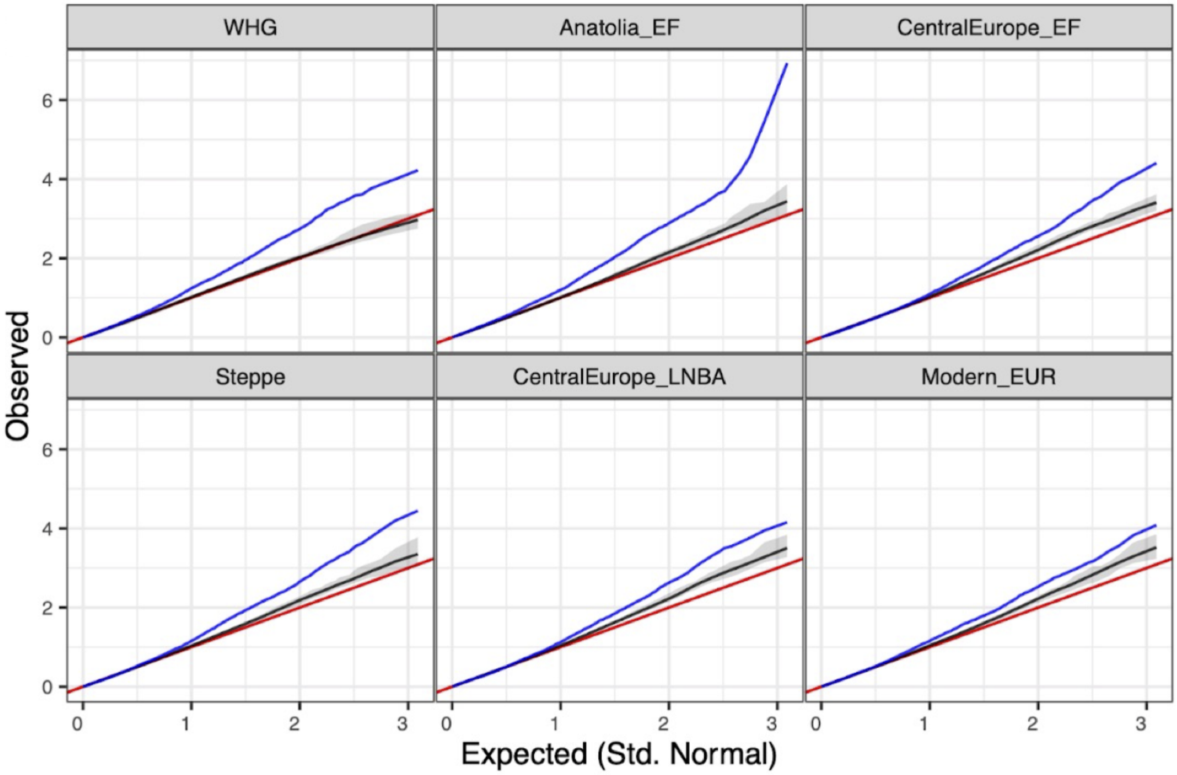


**Fig. S6.** QQ (quantile-quantile) plots of gene scores from empirical (blue line) and simulated (black line) genetic datasets for six different populations (note that only positive *Z* scores are plotted; see SI Appendix Methods). The red line shows the expected 1:1 relationship if gene scores exhibit a standard normal distribution. The results are consistent with the FPR analyses (Figs. 2B), with a slight excess of low *p*-values amongst all simulated populations other than WHG.

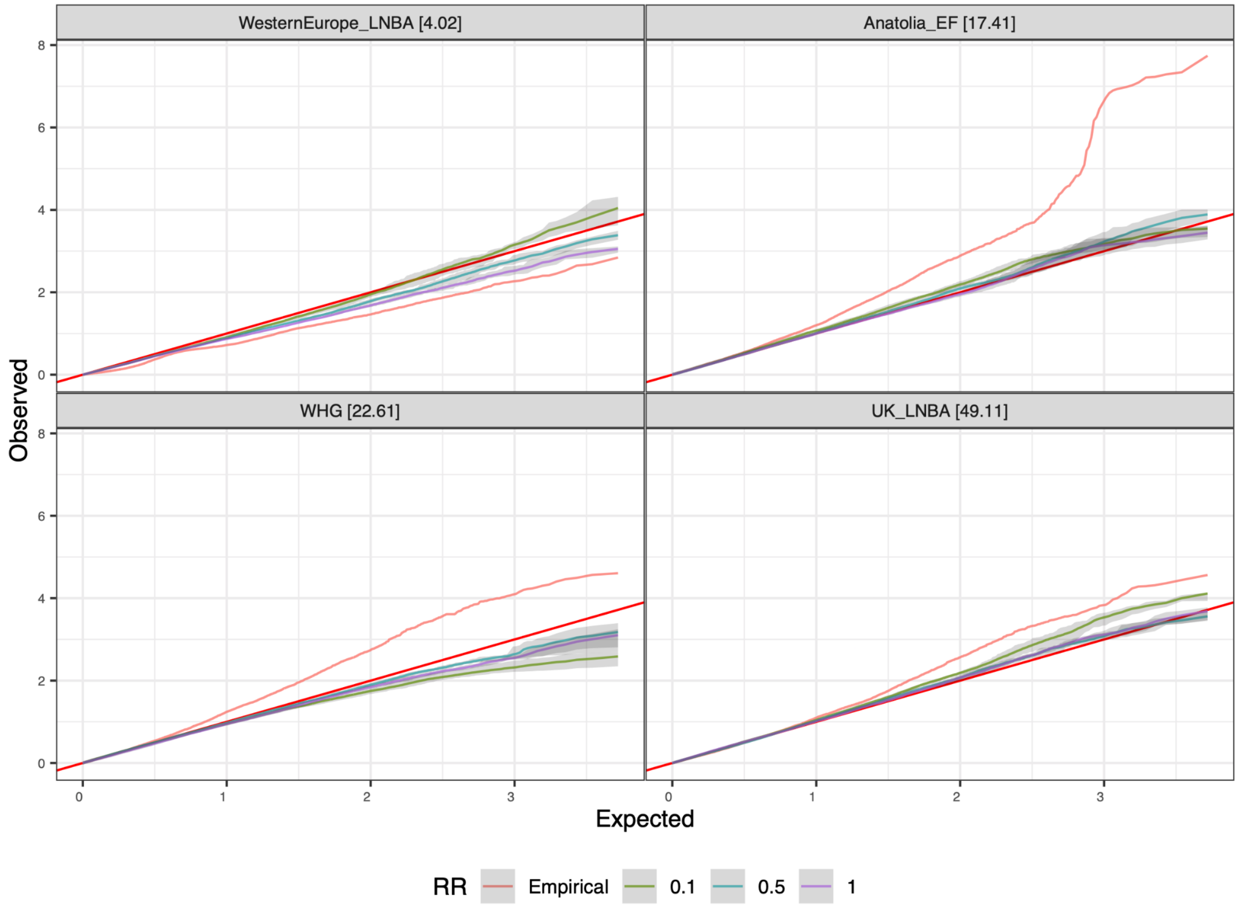


**Fig S7.** QQ (quantile-quantile) plots of gene scores from empirical (pink line) and simulated (black line) genetic datasets using different recombination rates (see key; factor of reduced recombination rate, assuming a default of 1.45 cM/Mbp), for four different populations (note that only positive z-scores are plotted; see SI Methods). The red line shows the expected 1:1 relationship if gene scores exhibit a standard normal distribution. Effective sample sizes (see Methods) are reported in square brackets next to each population name.

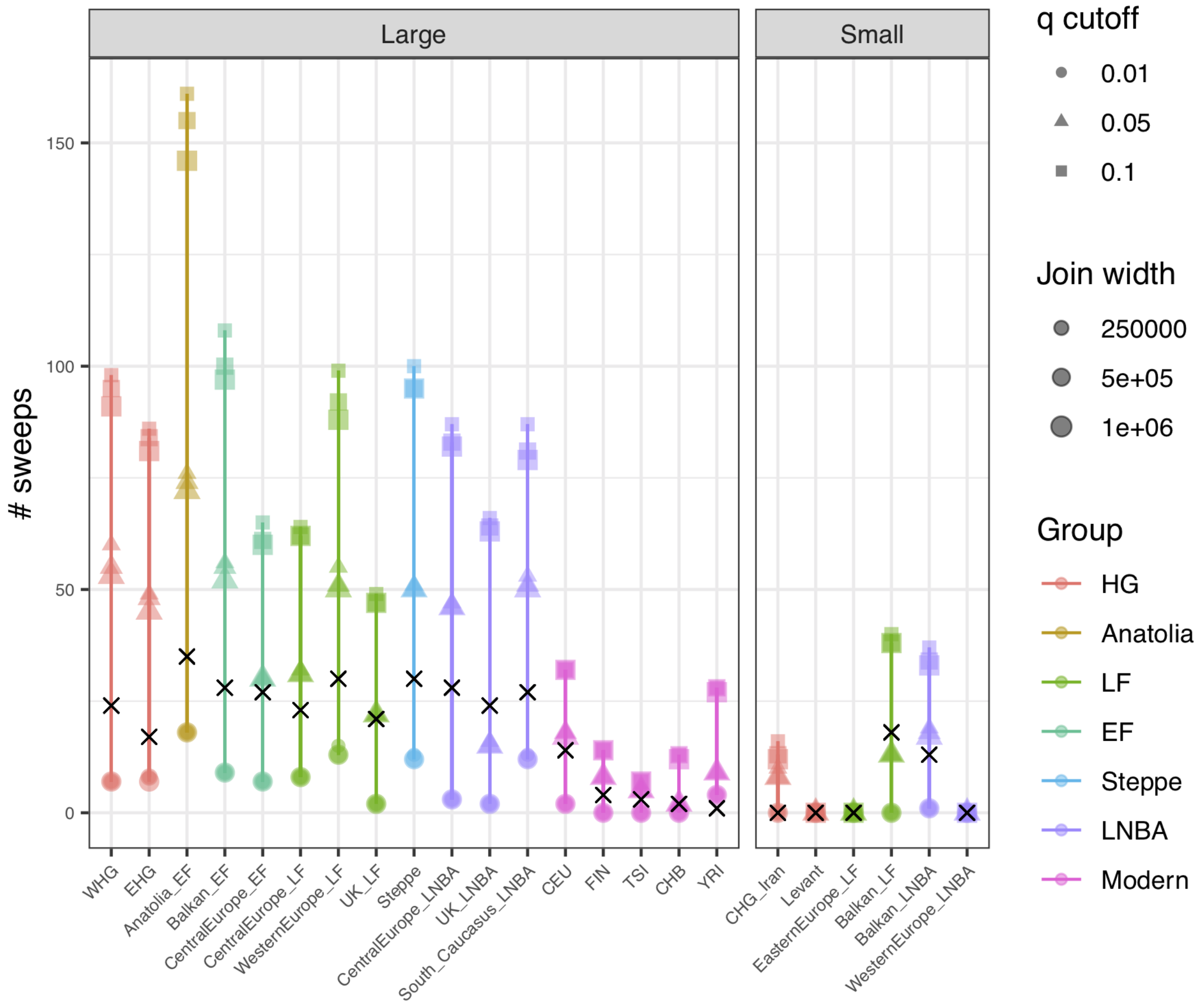


**Fig. S8.** The frequency of sweeps across populations according to different *q*-value thresholds and outlier gene aggregation distances. Sweeps were defined by grouping together outlier genes within 250kb, 500kb, or 1Mb (i.e. join-width) of each other that fell under a specific *q*-value threshold (see key). The sweep binning criterion used in the present study defined outlier genes at $q < 0.10$ and used a join-width of 1Mb to aggregate into sweeps. However, only sweeps with at least one outlier gene with $q < 0.01$ were retained as candidate sweeps. These sweeps are indicated by the 'x' symbol in the plot. Populations were split into two groups based on whether they had sufficient data to make robust inferences ($n_{eff} \geq 10$ = 'Large') or not ($n_{eff} < 10$ = 'Small') (see Methods).

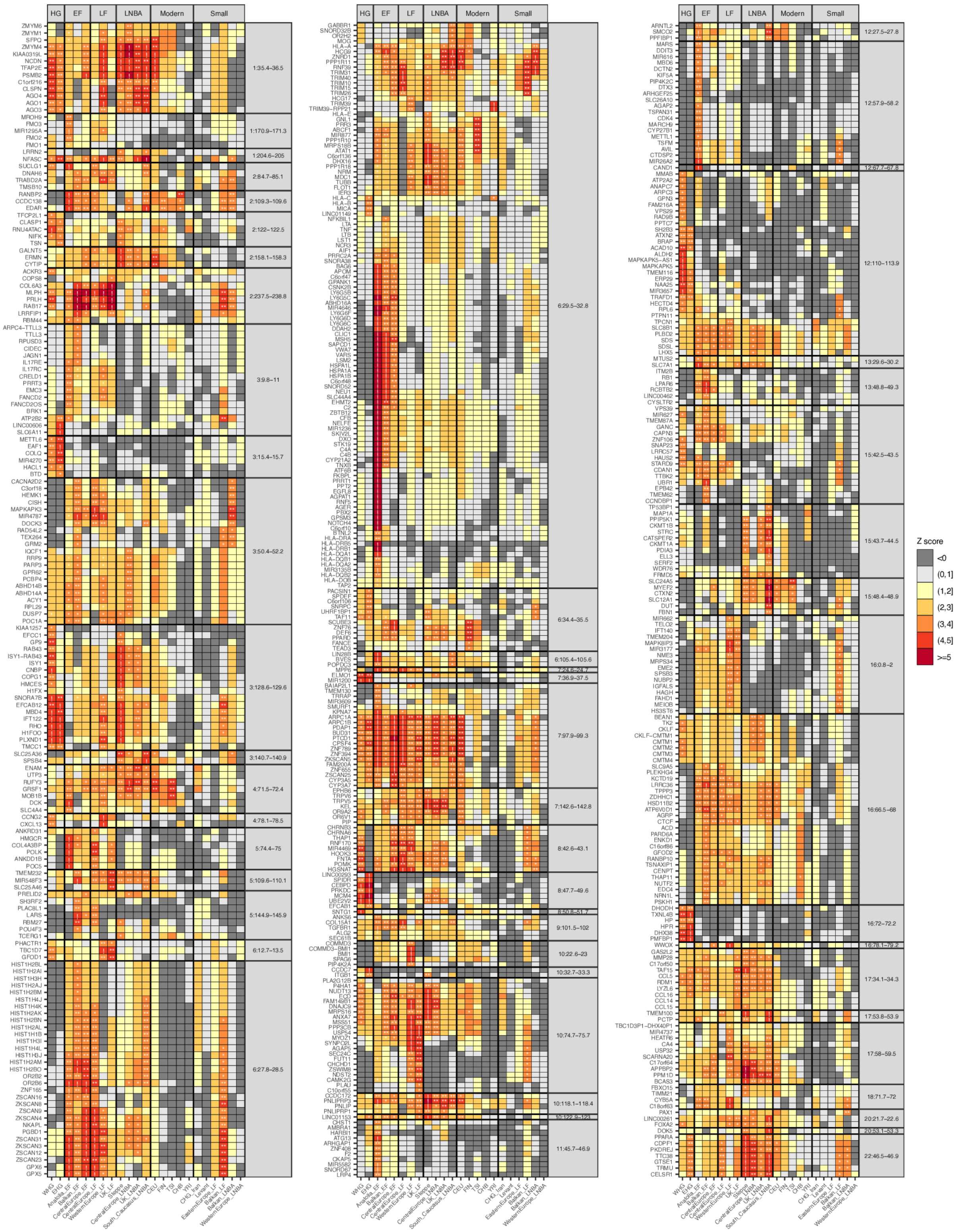
HG
EF
LF
LNBA
Modern
Small
Z score
<0
(0,1]
(1,2]
(2,3]
(3,4]
(4,5]
>=5

**Fig. S9.** The *Z*-score for each outlier gene observed amongst the 57 candidate sweeps (y-axis), for all modern and ancient populations (x-axis). Populations are organized chronologically according to broad archaeological categorizations, with populations that had insufficient data for robust inferences labelled as 'Small' (i.e. $n_{eff} < 10$; see Methods). Sweeps and genes are organized by chromosomal position. Outlier genes are indicated with a specific symbol according to their $q$-value ($0.05 \leq q < 0.10$ = *; $0.01 \leq q < 0.05$ = **; $q < 0.01$ = !).

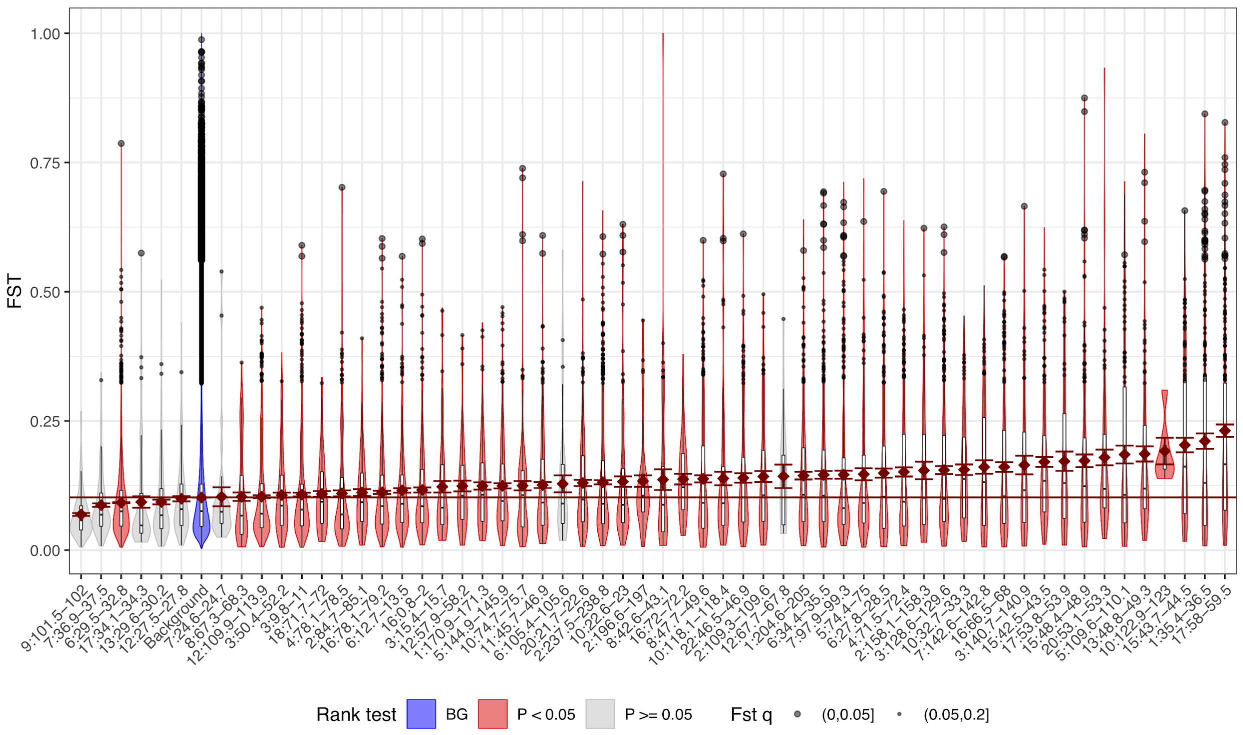


**Fig. S10.** Distribution of *F*st scores – evaluated for the 13 ancient western Eurasian populations with $n_{\text{eff}}$ ≥ 10 and a modern African population (YRI) – at each SNP within the 57 candidate sweeps (red or grey violins) vs. the remaining genomic background (blue violin). More than half (29/57) of the candidate sweeps contained significantly divergent SNPs ($q < 0.05$, large black dots; see (*6*)), and nearly all (55/57) contained strongly divergent SNPs ($q < 0.20$, small black dots). Further, 49 of the 57 sweeps had significantly higher *F*st scores than the remaining genomic background (Wilcoxon Rank Sum Test $p < 0.05$; significant = red violins; not significant = grey violins), consistent with elevated divergence in the candidate sweep regions due to positive selection.

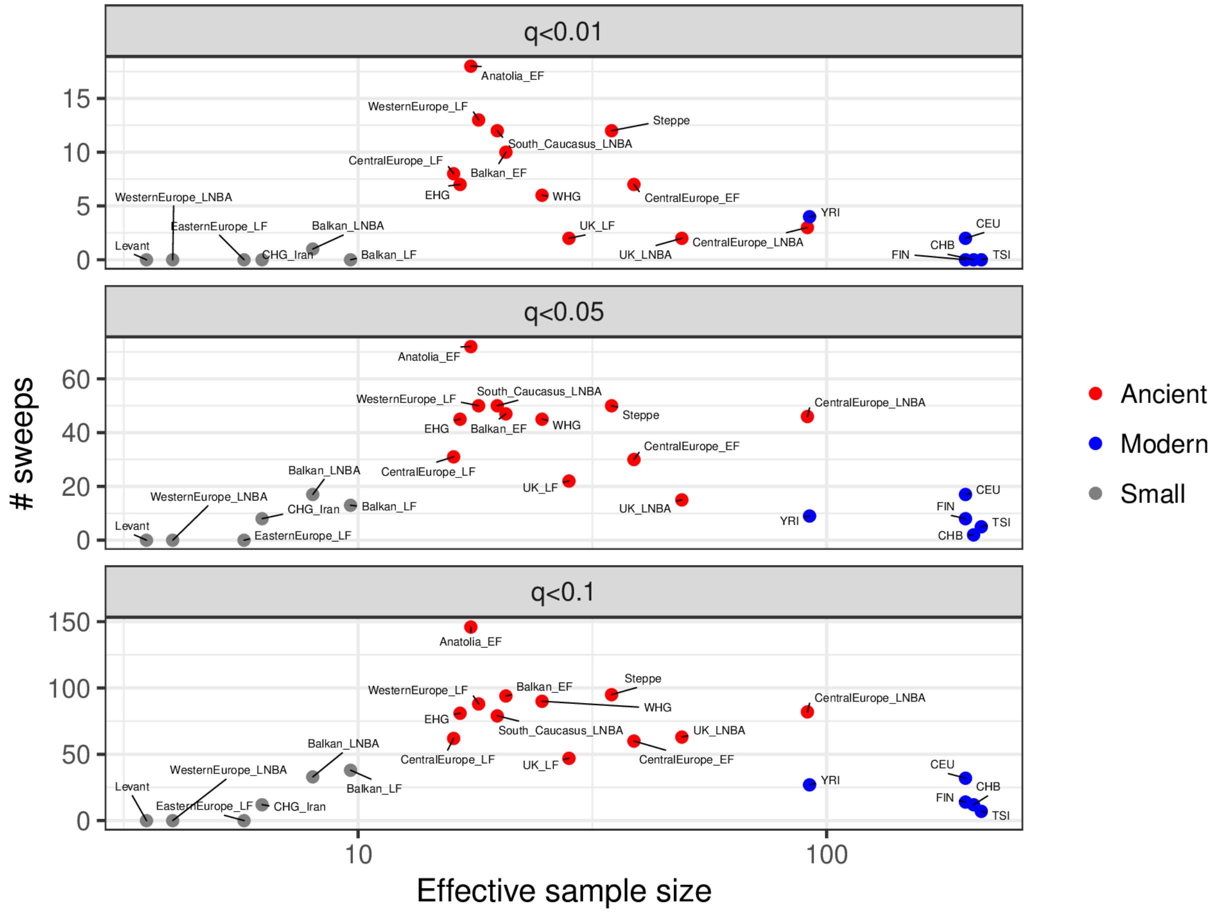


**Fig. S11.** The frequency of sweeps across populations relative to the effective sample size for each population. Horizontal panels show different *q*-values used to determine the outlier sweeps. Notable decreases in the number of observed sweeps occur for small populations (i.e. $n_{\text{eff}} < 10$; gray dots) and also modern European populations (blue dots), though only the former is likely to have been a consequence of low statistical power.

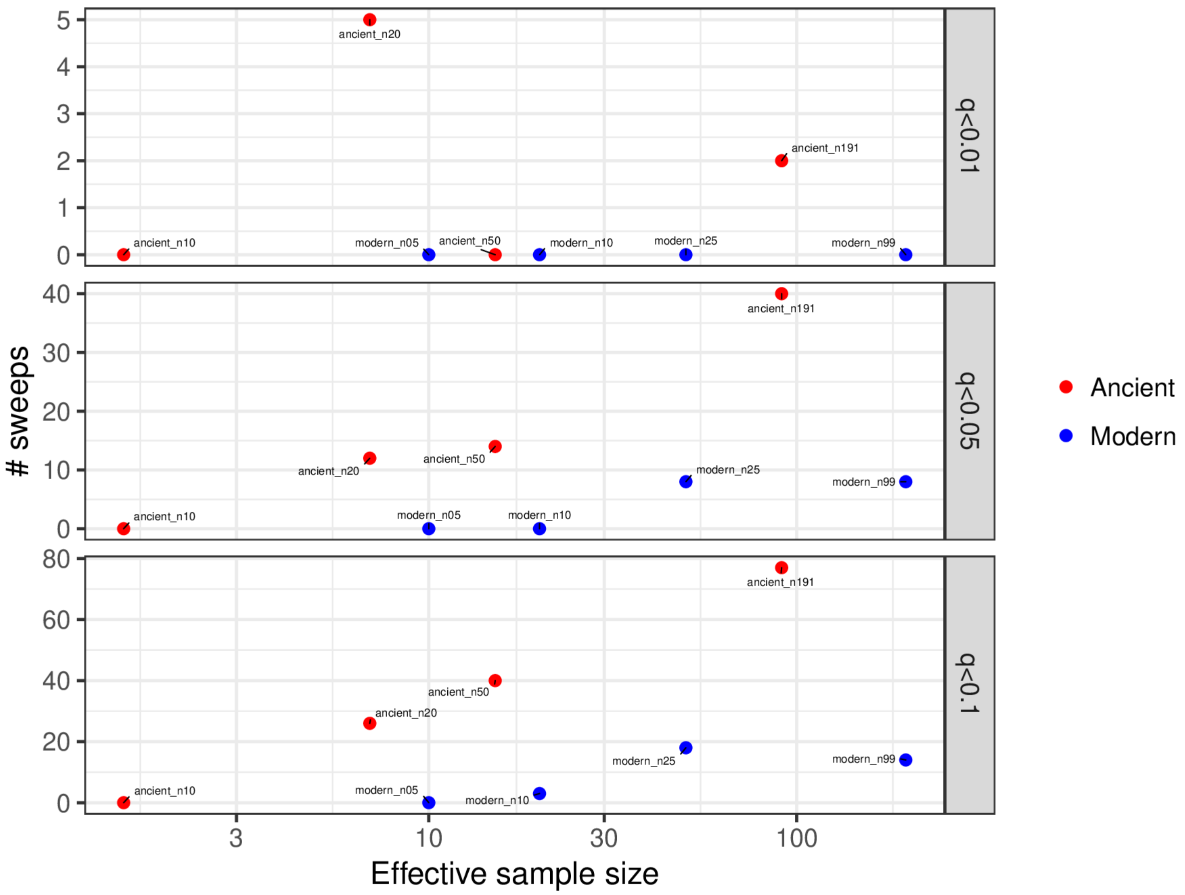


**Fig. S12.** The frequency of sweeps relative to effective sample size, $n_{\text{eff}}$, across an ancient (red dots) and a modern (blue dots) population resampled to smaller effective sample sizes (the number of samples per population, $n$, is indicated in labels next to each point, with the resulting $n_{\text{eff}}$ on x-axis). Horizontal panels show different $q$-values used to determine the outlier sweeps. There is a general tendency for the number of detected sweeps to decrease with smaller $n_{\text{eff}}$, particularly for less stringent $q$-value cutoffs where stochasticity is less likely to be a factor. However, the number of detected sweeps always remains smaller for the modern population relative to the ancient population for comparable $n_{\text{eff}}$ values.

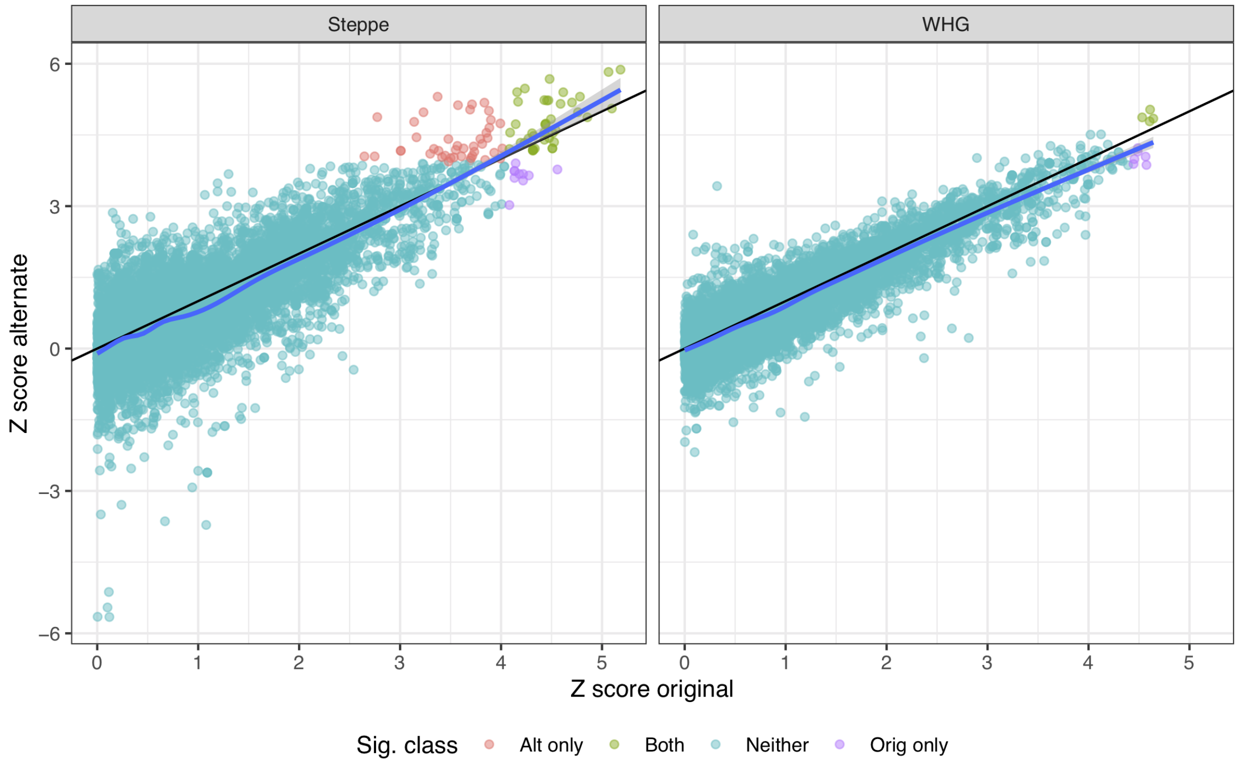


**Fig. S13.** Distribution of gene scores for two alternate sample groupings for the WHG and Steppe populations (see Table 1). The gene scores are strongly correlated (Pearson's $r = 0.82$ and 0.64 for *WHG* and *Steppe*, respectively), though outliers ($q < 0.01$) differ between the alternate sample groupings. Nonetheless, the shared ancestry amongst the ancient western Eurasian populations means that a similar set of outlier sweeps were observed using either groupings, as the sweep signals tend to be observed across multiple ancient populations (see Figs. S9, S14).

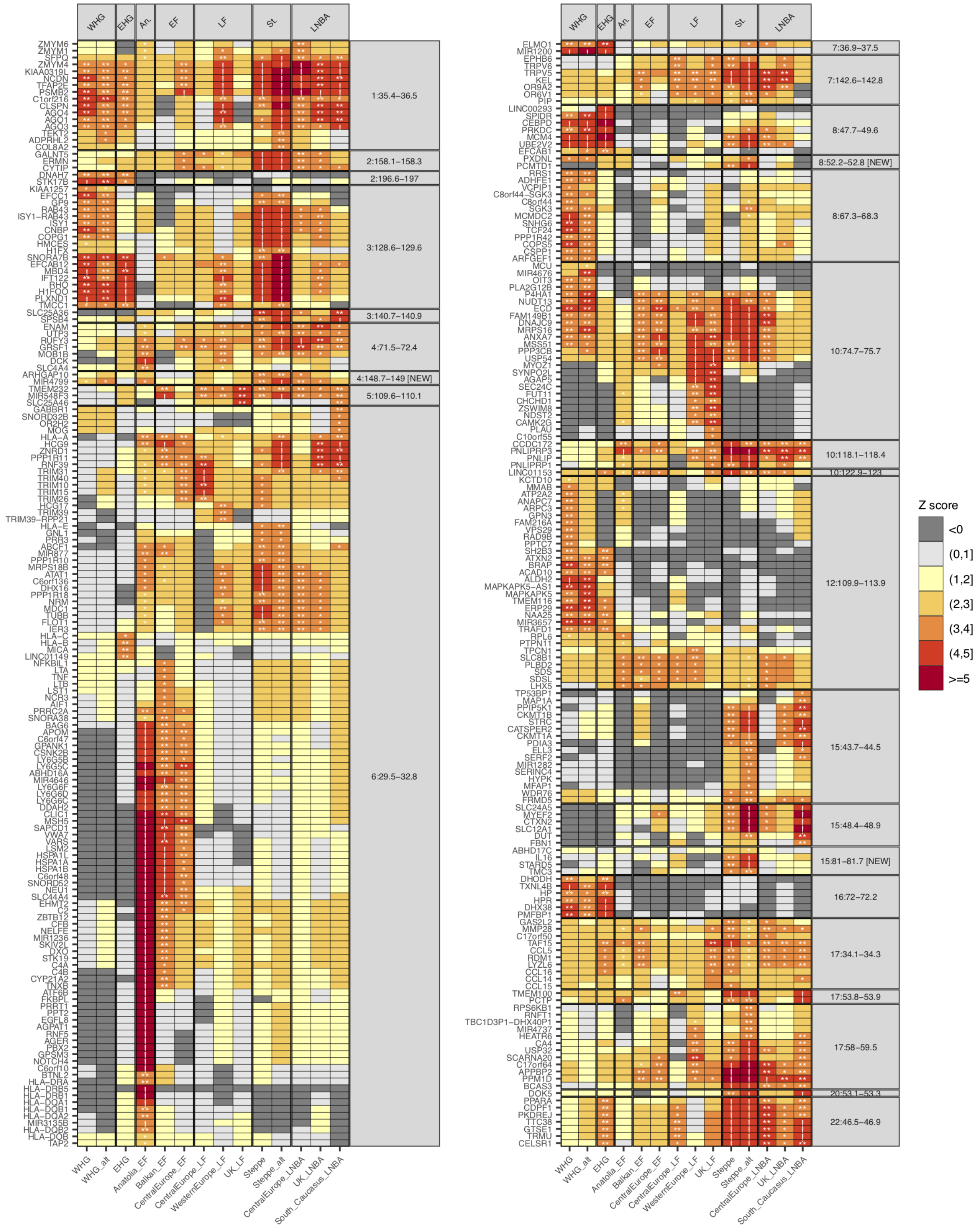

WHG
EHG
An.
EF
LF
St.
LNBA
1:35.4–36.5
2:158.1–158.3
2:196.6–197
3:128.6–129.6
3:140.7–140.9
4:71.5–72.4
4:148.7–149 [NEW]
5:109.6–110.1
6:29.5–32.8
7:36.9–37.5
7:142.6–142.8
8:47.7–49.6
8:52.2–52.8 [NEW]
8:67.3–68.3
10:74.7–75.7
10:118.1–118.4
10:122.9–123
12:109.9–113.9
15:43.7–44.5
15:48.4–48.9
15:81–81.7 [NEW]
16:72–72.2
17:34.1–34.3
17:53.8–53.9
17:58–59.5
20:53.1–53.3
22:46.5–46.9
WHG
WHG_alt
EHG
Anatolia_EF
Balkan_EF
CentralEurope_EF
CentralEurope_LF
WesternEurope_LF
UK_LF
Steppe
Steppe_alt
CentralEurope_LNBA
UK_LNBA
South_Caucasus_LNBA
Z score
<0
(0,1]
(1,2]
(2,3]
(3,4]
(4,5]
>=5

**Fig. S14.** The *Z*-score for each gene amongst the 27 sweeps (y-axis) observed for *WHG* and *Steppe* populations, using two different sample groupings for each population (see SI Methods). Sweeps and genes are organized by chromosomal position. Outlier genes are indicated with a specific symbol according to their $q$-value ($0.05 \le q < 0.10$ = *; $0.01 \le q < 0.05$ = **; $q < 0.01$ = !). Importantly, while changing the sample grouping of the *WHG* and *Steppe* populations resulted in some different sweeps being detected for each, 19 of the 27 sweeps were observed regardless of which sample grouping was used – 12 were observed in both sample groupings and the remaining seven were observed in one or more population other than the *WHG* or *Steppe*. Our results imply that the shared genetic history and large number of ancient populations used in this study increase the chance that a sweep will be observed and thereby improve the robustness of the sweep detection process.

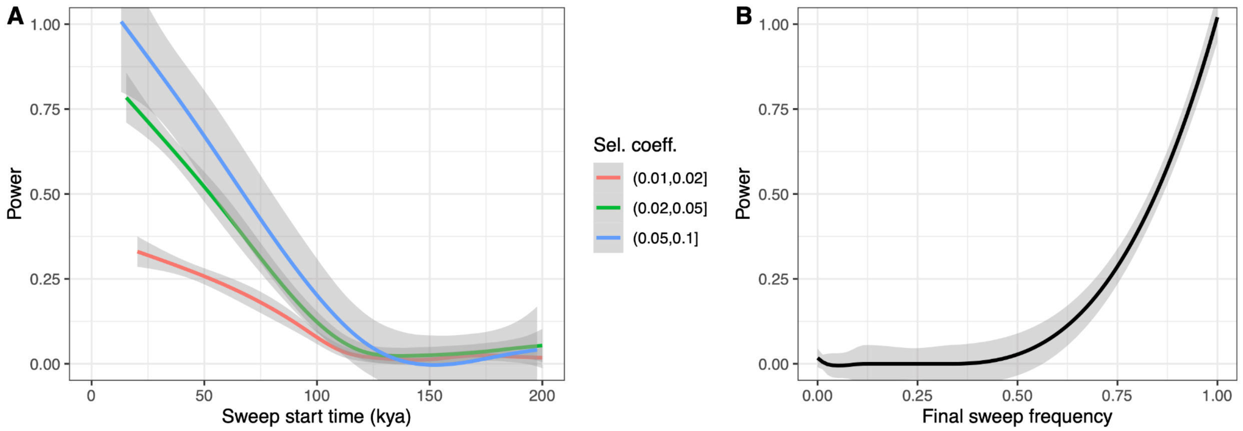


**Fig S15.** Detection power for sweeps based on (A) selection start time and (B) final frequency at the time of genome sampling. Selection was simulated with coefficients ranging between 1% and 10% within a recently published population demography[11], with genomic sampling performed in the Anatolian EF population at 8ka. In total, 10,000 sweeps were simulated with start times uniformly distributed between 8ka and 200ka. Admixture from European hunter-gatherers, Steppe, Basal Eurasian or Neanderthal populations was excluded from the demography to explicitly quantify power in the absence of admixture. In (A), only sweeps with a final frequency of more than 80% were considered, whereas (B) includes incomplete sweeps that started less than 20kyrs prior to sampling. Power falls below 50% when selection started prior to ~80ka (i.e. ~70ka before sampling occurred), or when the final haplotype frequency is below 85% at the time of sampling.

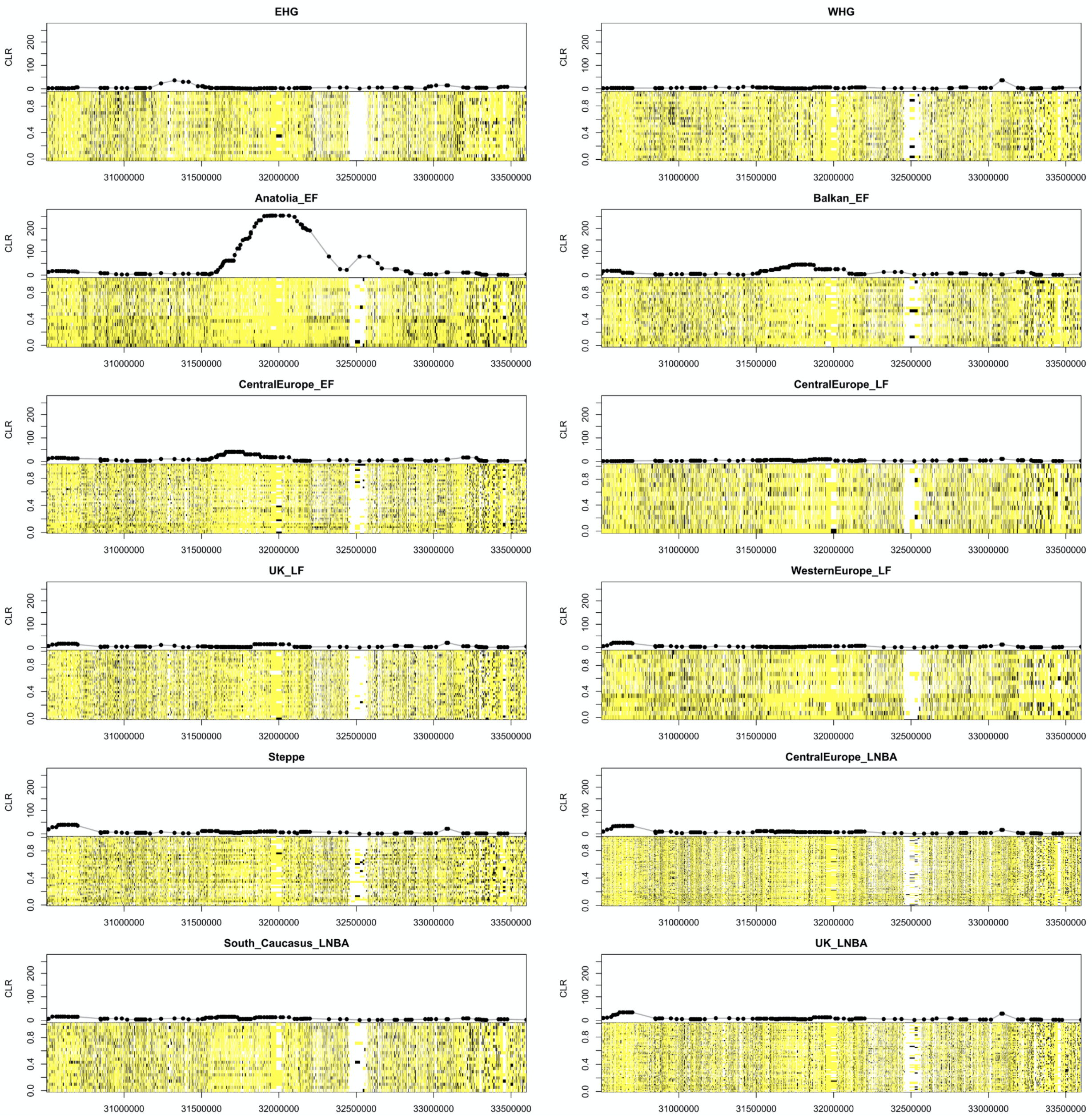


**Fig S16.** Haploimage of the MHC-III region for all ancient populations. Pseudohaplotype calls are shown for all samples in each population, with majority alleles in yellow, minority alleles in black, and missing data in white. The SweepFinder2 CLR scores for all annotated genes within the sweep region are shown on top of each haploimage. Haploimages for all sweeps are provided as Supplementary Data 58-114.

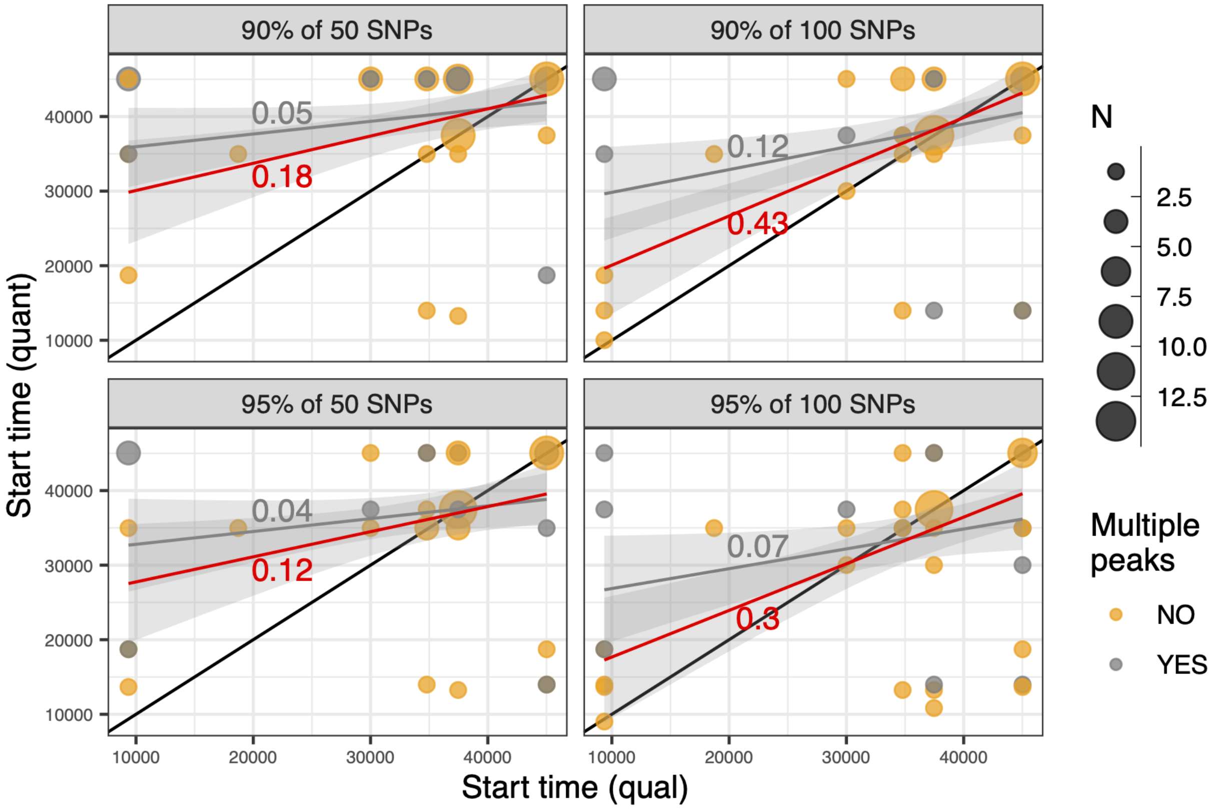


**Fig S17.** Comparison between qualitative (x-axis) and quantitative (y-axis) sweep haplotype detection methods. Sweeps were assigned an onset time by scanning five Upper Paleolithic specimens for evidence of the sweep haplotype, with the oldest sample in which the haplotype was observed serving as a coarse lower bound for the origin of the sweep (see Methods). Results are shown for the four different combinations of the two detection criteria used for our quantitative method (see SI Methods). Pearson's correlation coefficient, *r*, was weakly positive when measured on all sweeps (gray lines, *r* shown above line; sweep counts indicated by gray circles, see key), but improved markedly when excluding the 12 sweeps that had multiple SF2 peaks (red lines, *r* shown below line; sweep counts indicated by orange circles).

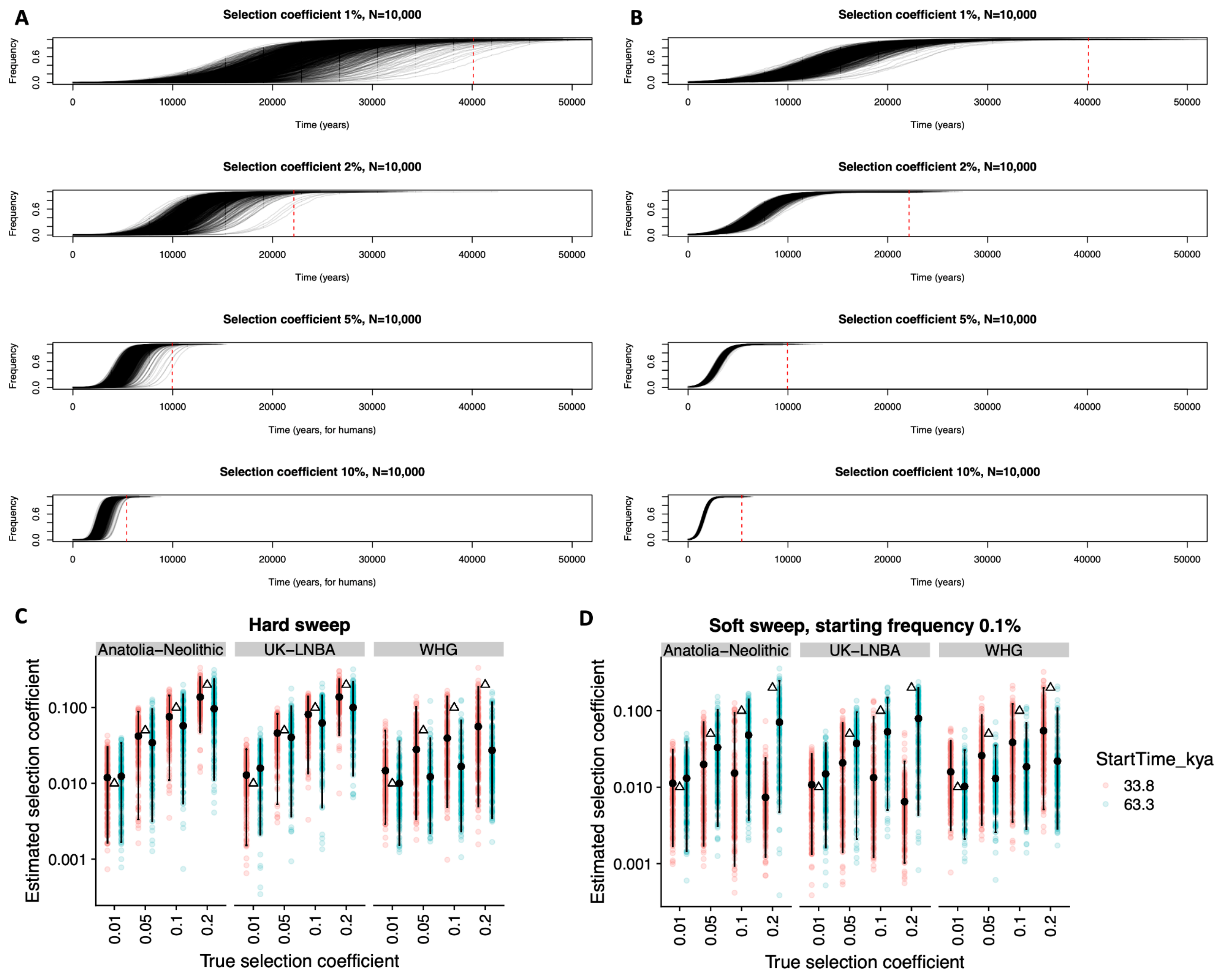


**Fig. S18.** Estimating selection strength and fixation time properties of the 57 sweeps. (A) Time to fixation for a beneficial *de novo* mutation for 1000 forward-in-time simulations at each of four different selection coefficients, *s*, ranging between 1% and 10% (see SI Methods). The beneficial *de novo* mutation arises at time zero and the simulations are conditioned on the fixation of this allele. We assume a generation time of 30 years and a single randomly mating population of 10,000 diploid individuals. The red dashed line indicates the expected time to fixation for a beneficial *de novo* mutation using an analytical formula from ref. [37]. (B) Same as panel A, but the beneficial allele starts at a frequency of 0.1% at time zero. (C) Estimation of the selection coefficient, *s*, for simulated sweeps from *de novo* mutations and (D) rare standing variation (0.1% frequency). Simulations were performed with two different selection start times (see key). The simulated demography is taken from ref. [11] but excludes any admixture events. The sample sizes and missing data distributions are taken from the respective empirical data. The simulated *s* is shown as a triangle and the inferred values depicted as coloured circles (see Methods). Estimated *s* values behave reasonably well for simulated *s* = 1%, though tend to be systematically downwardly biased for *s* values of 5% and larger – particularly for recently selected standing variants – suggesting that the selection coefficients reported for most of the 57 candidate sweeps are also probably underestimated.

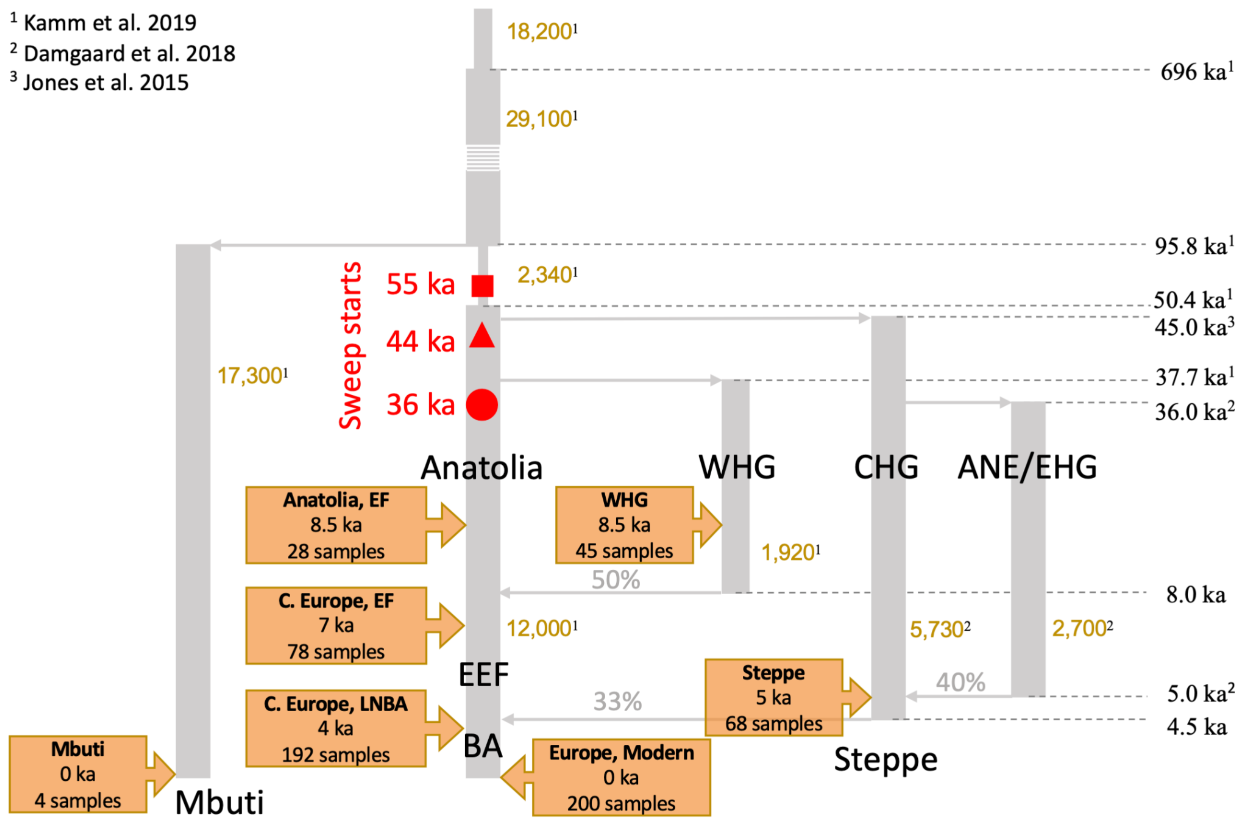


**Fig. S19.** Schematic of the West Eurasian population history model used to explore the statistical properties of our analytical pipeline. Each vertical segment denotes a major population branch (effective population sizes shown in gold text), with grey horizontal arrows denoting separation and admixture events (times shown on the right-hand side of the figure; percentages indicating the proportion of ancestry contributed by the incoming admixture branch). Model parameters are taken from one of three studies, as denoted by the associated superscript (1 = ref. [11]; 2 = ref. [41]; 3 = ref. [42]). Hard sweeps were simulated on the Main Eurasian branch at three different times (red shapes and text) and were inherited by all descendant populations. The yellow text boxes provide information on the samples that were used in simulation analyses (haploid sample sizes reported), which occur before and after major admixture events.

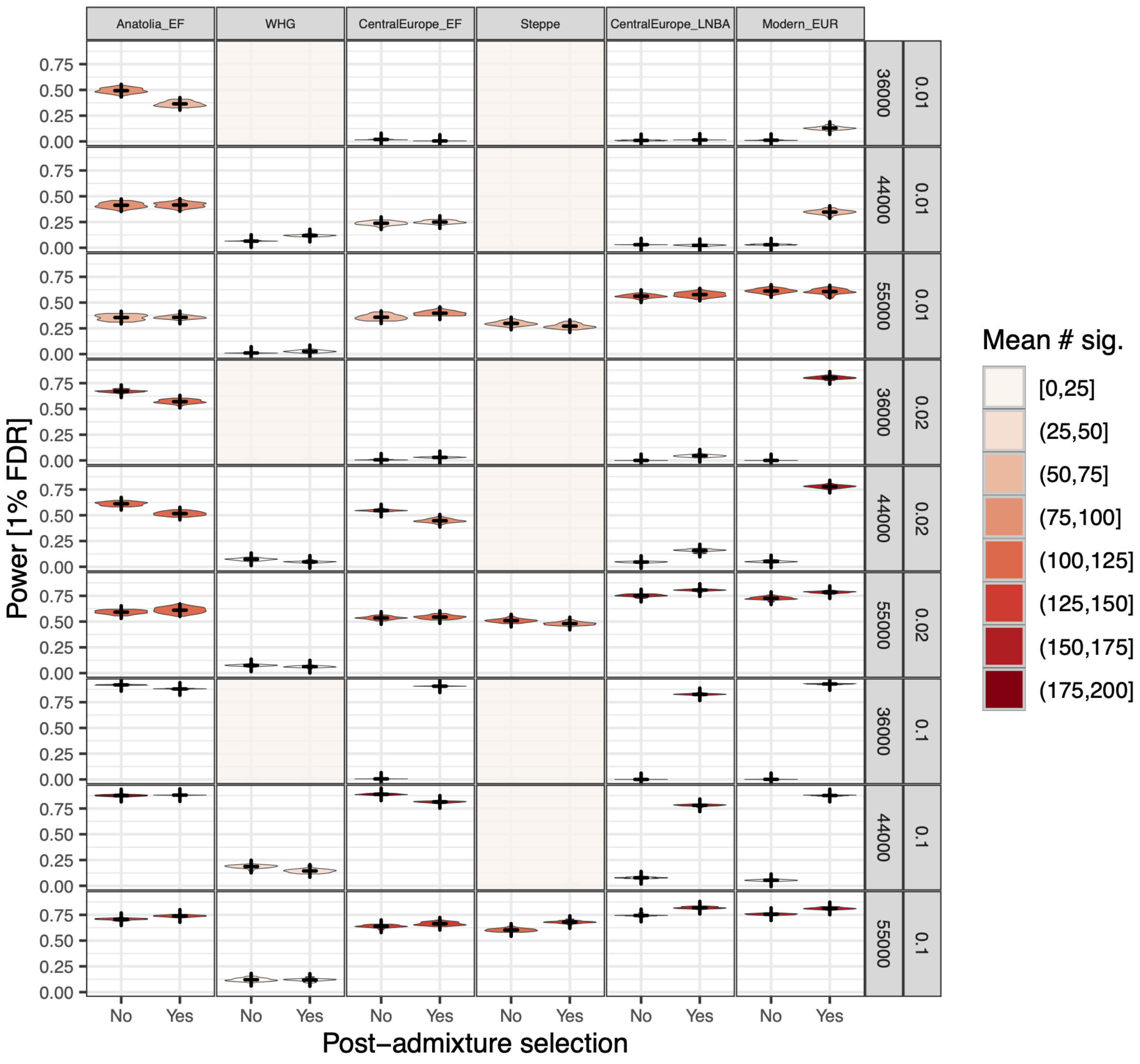


**Fig. S20.** Estimated power of the analytical pipeline. Results are plotted for 200 simulated genomes for each of six simulated populations (column panels) and all combinations of selection strength (row panels, see outer labels) and starting time (row panels, see inner labels). Each violin is coloured according to the number of significant tests, with mean and standard deviations indicated by the black horizontal and vertical lines, respectively. Selection was either modelled as acting uniformly across the entire population history (x-axis, post admixture selection = No), or ceasing at 8ka following the admixture of WHG and Main Eurasian branch (post admixture selection = Yes). Panels with no information represent populations that split from the Main Eurasian branch before the introduction of the beneficial allele that therefore did not inherit the selected locus.

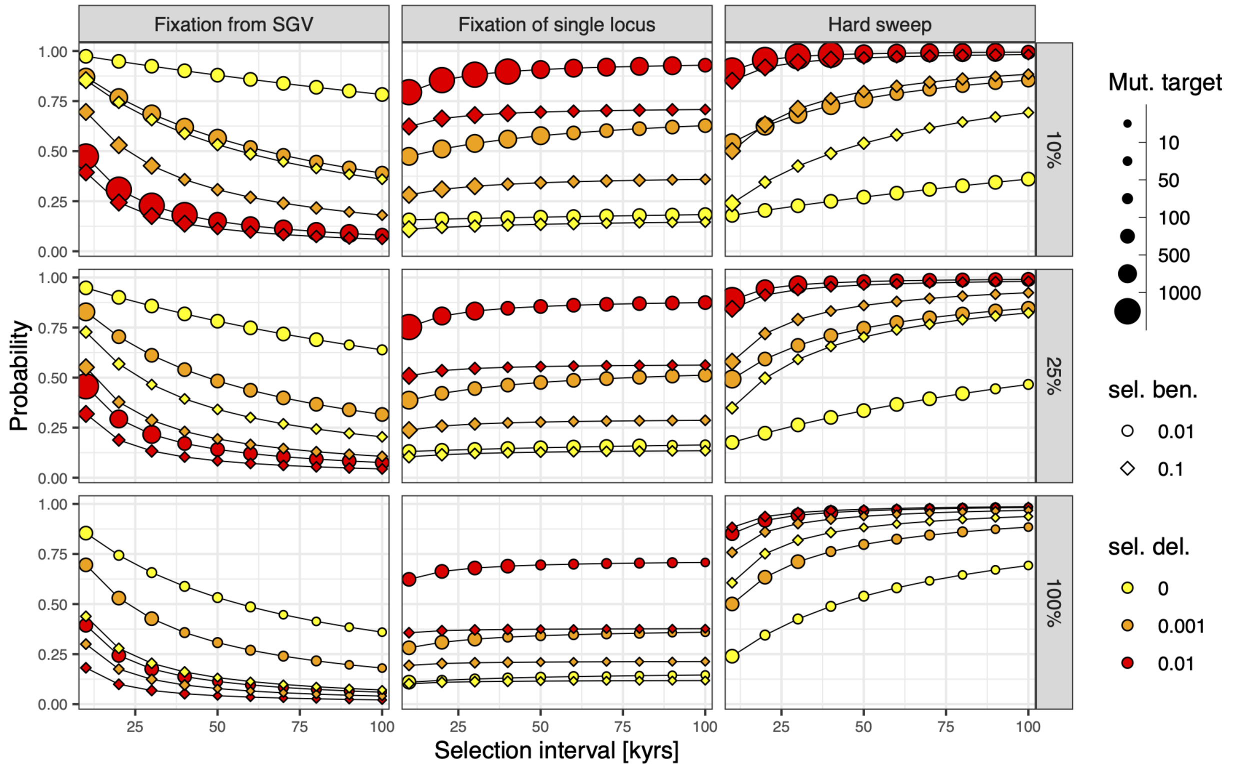


**Fig. S21.** The probability that a sweep was caused by a standing genetic variant (SGV) relative to a *de novo* mutation conditional on a sweep of either type occurring within a prescribed time interval (left panels). Symbol sizes show the expected mutational target sizes needed to ensure fixation of the beneficial allele within a specific time interval (indicated on the x-axis). SGVs are assumed to have been under some degree of purifying selection (denoted by different coloured symbols) prior to the environmental shift that initiates the beneficial selection phase (symbol shapes), with Eurasian populations experiencing a range of bottleneck intensities (Eurasian $N_e$ relative ancestral $N_e$ indicated in row panels). The probability that all fixed SGVs descended from a single copy at the time of the environmental shift (central panels) was strongly dependent on the strength of purifying selection, reversing the direction of this dependency observed for the SGV fixation probabilities. Combining these two probabilities resulted in hard sweep patterns from either SGVs or *de novo* variants being highly likely (>50%) when causal mutations were at least weakly deleterious prior to the environmental change (right panel; see Text S5). All equations come from ref. [37].

#### Supplementary Tables

**Table S1.** Full set of metadata for all ancient western Eurasian samples considered in this study. Data includes sample ages, geographic locations, as well as associated cultural and historical affinities for each individual. This information is combined with genetic relationships between samples in order to categorize the populations that were used for selection analyses.

**Table S2.** List of 57 outlier sweeps at $q$-value < 0.01. Sweeps are labelled according to their chromosome position, with full positional details shown in subsequent columns. Each sweep is dated according to its first appearance amongst five Paleolithic individuals in our dataset (see Methods). Selection results are provided from two previous studies and were used as a comparative dataset. Selection coefficients for each sweep were estimated using SweepFinder2 and local recombination rate (see Methods). The full list of genes present in each sweep are given in the last column.

**Table S3.** The number and proportion of mappable SNPs falling within each of the 57 candidate sweep windows relative to other regions of the genome, based on the CRG100 mappability scores from ref. [15]. All sweeps contain at least 96.7% mappable SNPs, indicating that they are unlikely to be alignment artefacts.

**Table S4.** Parameter values used to calculate the probability of fixation from standing genetic variation and *de novo* mutations, based on equations from ref. [37].

#### Supplementary Data

**Supplementary Data 1-57**. Composite likelihood ratio (CLR) scores (grey line) as a function of genomic coordinates (x-axis) for the top 57 candidate sweeps. Genes within the sweep region are represented by colored boxes, with the color intensity indicating their Z-scores. For each sweep, up to 10 genes with $q$-values less than 0.01 are named. If more than 10 genes have $q$-values < 0.01, then the 10 genes with the highest aggregate $Z$ score across all populations are reported.

The SNP marker weights used to determine the selected haplotype for each sweep are shown in the top panel, with the SNPs used for the top 30, 50, and 100 markers indicated in the key.

**Supplementary Data 58-114**. HaploImages for each of the 57 candidate sweeps (*q*-value < 0.01). The images depict the pseudo-haplotype calls at each locus in the sweep window and flanking region, for each population used in this study. Majority alleles, *i.e.* those alleles most likely to be associated with the sweep, are shown in yellow, and minority alleles in black, with missing data being white. Only individuals with no more than 50% missing data are shown. In addition to the populations scanned for sweeps using SweepFinder2, haplo-images for five Late Pleistocene Eurasian AMH samples (Ust'-Ishim, Kostenki14, GoyetQ-116, and El Mirón) were also included to help determine the earliest timing of the selection pressure for each sweep. Note that only Ust'-Ishim had sufficient coverage to make diploid calls (*i.e.* both alleles are present at each locus), whereas the four remaining samples use pseudo-haploid calls.
